# A Cholinergic Signaling Pathway underlying Cortical Circuit Regulation of Lateral Ventricle Quiescent Neural Stem Cells

**DOI:** 10.1101/2023.09.02.556037

**Authors:** Moawiah M Naffaa, Henry H. Yin

## Abstract

Neurogenesis and proliferation of neural stem cells (NSCs) in the subventricular zone (SVZ) are controlled by both intrinsic molecular pathways and extrinsic signaling cues, including neural circuits. One such circuit, the ACC-subep-ChAT^+^ circuit, has been identified as a regulator of ventral SVZ neurogenesis by modulating the proliferation of LV NSCs. However, the specific neural signals that promote the proliferation activity of LV NSCs have remained largely unknown. In this study, we uncover a molecular mechanism underlying the cellular activation and proliferation of quiescent NSCs (qNSCs) in the lateral ventricle SVZ (LV-SVZ) mediated by the cortical circuit. Our findings demonstrate that postnatal and adult LV qNSCs are triggered by the cortical circuit through ChAT^+^ neuron stimulation, consequently resulting in the activation of muscarinic 3 receptors (M3) expressed on LV qNSCs. This, in turn, triggers inositol 1,4,5-trisphosphate receptor type 1 (IP3R1) activation, causing intracellular calcium release. Subsequently, the proliferation of LV qNSCs occurs through the downstream regulation of the calcium/calmodulin dependent protein kinase II delta (CAMK2D) and the MAPK10 signaling pathway. These findings shed light on the molecular regulatory mechanisms that govern LV qNSCs and emphasize the significant role of the cortical circuit in promoting their proliferative activation within the ventral LV-SVZ.

## Introduction

Postnatal and adult mammalian neurogenesis occurs in the subventricular zone (SVZ) of the lateral ventricles (LV) (1). LV neural stem cells (LV NSCs) could be quiescent or actively dividing (2, 3). Upon activation, quiescent LV NSCs can divide asymmetrically for self-renewal or differentiate into transient amplifying intermediate progenitors (TAPs) (2, 4, 5). These progenitors undergo symmetrical division and differentiation, producing young migrating doublecortin-positive (DCX^+^) neuroblasts (6, 7). The neuroblasts migrate to the olfactory bulb (OB) (6, 8, 9), where they become mature interneurons which are integrated into the local circuitry (10, 11). Adult-born young neurons from the LV niche plays a crucial role in experience-dependent plasticity in the postnatal and adult brain (12, 13), particularly for olfaction-based behaviors in rodent (11, 14-16). Elucidating the regulatory mechanisms of SVZ neurogenesis including LV qNSCs activation and proliferation is critical to our understanding of neuronal regeneration (17, 18), neurological disorders, and brain injury (19, 20).

Numerous intrinsic and extrinsic factors control the renewal and differentiation of LV NSCs, distinct from embryonic NSCs (21, 22). As an example of regulatory factors, LV NSCs receive signals from both local and distant neurons through neurotransmitter release (22). They respond to various neurotransmitter signals that regulate postnatal and adult SVZ neurogenesis (23-25). Yet the mechanisms governing the control of LV NSC proliferation activity by various neurotransmitters remain inadequately studied. An example of these neuronal regulators is subep-ChAT^+^ neurons, which release acetylcholine (ACh) locally in the ventral SVZ and modulate LV NSC proliferation and neuroblasts generation (26). These cholinergic neurons receive inhibitory inputs from local GABAergic neurons and excitatory inputs from the anterior cingulate cortex (ACC) (27). However, the molecular mechanisms underlying the circuit regulation of LV NSC proliferation remain unknown. Understanding these mechanisms could open a new avenue of research to explore how external cues regulate olfactory perception activity through cortical regulation of SVZ-olfactory neurogenesis.

In this study, we hypothesized the existence of a muscarinic regulatory pathway underlying the cortical regulation of the proliferative activity of LV qNSCs in the ventral SVZ. By employing functional, molecular, and *in vivo* neural modulation strategies, we demonstrated the role of muscarinic 3 receptors (M3) in regulating the activation of ventral LV qNSCs following ACC-subep-ChAT^+^ circuit activation. We have identified inositol 1,4,5-trisphosphate receptor type 1 (IP3R1) as a key component of the signaling pathway following M3 activation. This activation triggers intracellular calcium release, which in turn facilitates calcium/calmodulin-dependent protein kinase II delta (CAMK2D) signaling, and in initiates a downstream pathway involving Mitogen-activated protein kinase 10 (MAPK10), ultimately leading to LV qNSCs proliferation.

These findings offer insights into the molecular mechanism by which a cortical circuit initiates the activation and proliferation of LV qNSCs. This knowledge will contribute to future studies that investigate the regulation of neurogenesis within the SVZ through the activity of cortical neural circuits, as well as its relationship to learning behavior. Specifically, this molecular mechanism could be essential to understanding the role of the ACC circuit during postnatal development and associated disorders, such as mental disabilities.

## Results

### Activation and proliferation of LV qNSCs in ventral SVZ initiated by ACC-subep-ChAT^+^ circuit modulation

Our previous studies have demonstrated that stimulating the ACC-subep-ChAT^+^ circuit for a day leads to an increased proliferation of LV NSCs in close proximity to the subep-ChAT^+^ neurons (27). In order to investigate the molecular mechanism responsible for the circuit’s impact on cellular proliferation, we first used optogenetics to directly manipulate subep-ChAT^+^ neurons in this circuit for 6 hours only. To excite and inhibit subep-ChAT^+^ neurons, we used *ChAT-ChR2-EYFP* and *ChAT-Cre; AiD40* mice, respectively **(Fig. 1A and E)**. To detect proliferative cells, EdU was administered via intraperitoneal (IP) injection two hours before the end of the experiment.

**Figure 1.**
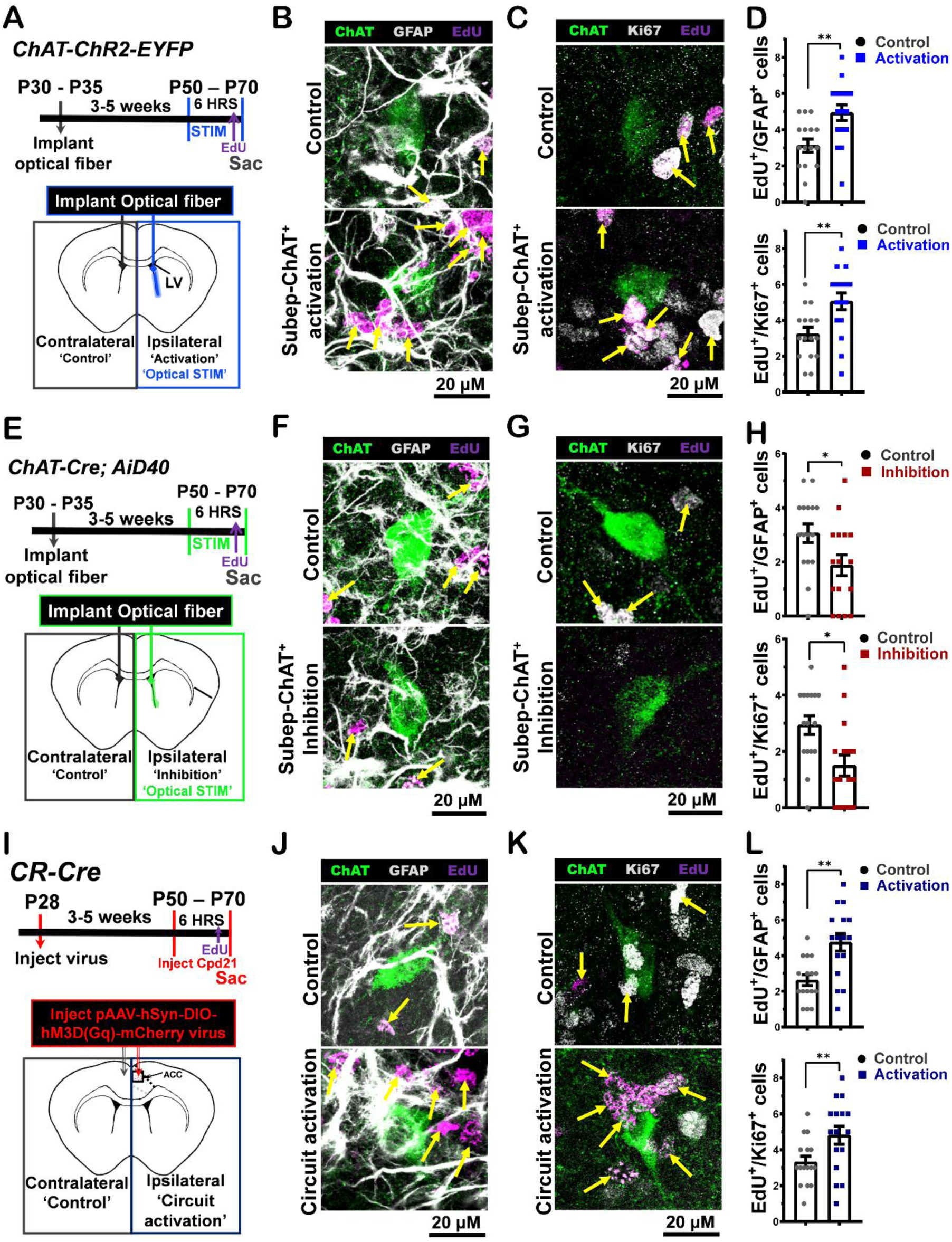
*In vivo* regulation of ventral LV NSCs via subep-ChAT^+^ neurons and ACC-subep-ChAT^+^ circuit modulation. **(A)** Experimental design (upper) and schematic representation (lower) of *in vivo* optogenetic activation for 6hrs post optical fibers implantation into LV regions of (P30) *ChAT-ChR2-EYFP* mice. **(B)** Immunofluorescence staining for ChAT (green), GFAP (gray) and EdU (purple) of ipsilateral SVZ wholemount (lower images; subep-ChAT^+^ neurons activation) vs. contralateral SVZ wholemount (upper images; (control)) from mice in panel A. Yellow arrows show EdU^+^/GFAP^+^ cells surrounding subep-ChAT^+^ neurons. Scale bar, 20 μm. **(C)** Immunofluorescence staining for ChAT (green), Ki67 (gray) and EdU (purple) of ipsilateral SVZ wholemount (lower images; subep-ChAT^+^ neurons activation) vs. contralateral SVZ wholemount (upper images; (control)) from mice in panel A. Yellow arrows show EdU^+^/Ki67^+^ cells surrounding subep-ChAT^+^ neurons. Scale bar, 20 μm. **(D)** Upper; analysis of EdU^+^/GFAP^+^ cells per a subep-ChAT^+^ neurons in SVZ wholemounts of ipsilateral vs. contralateral in panel B. P < 0.0017, t^15^ = 3.8, n=16, Paired t-test. Data collected from four stimulated *ChAT-ChR2-EYFP* mice. Each dot represents total EdU^+^/GFAP^+^ cells surrounding a subep-ChAT+ neuron. Lower; analysis of EdU^+^/Ki67^+^ cells per a subep-ChAT^+^ neurons in SVZ wholemounts of ipsilateral vs. contralateral in panel B. P < 0.0016, t^15^ = 3.9, n=16, Paired t-test. Data collected from four stimulated *ChAT-ChR2-EYFP* mice. Each dot represents total EdU^+^/Ki67^+^ cells surrounding a subep-ChAT+ neuron. **(E)** Experimental design (upper) and schematic representation (lower) of *in vivo* optogenetic activation for 6hrs post optical fibers implantation into LV regions of (P30) *ChAT-Cre; AiD40* mice. **(F)** Immunofluorescence staining for ChAT (green), GFAP (gray) and EdU (purple) of ipsilateral SVZ wholemount (lower images; subep-ChAT^+^ neurons inhibition) vs. contralateral SVZ wholemount (upper images; (control)) from mice in panel E. Yellow arrows show EdU^+^/GFAP^+^ cells surrounding subep-ChAT^+^ neurons. Scale bar, 20 μm. **(G)** Immunofluorescence staining for ChAT (green), Ki67 (gray) and EdU (purple) of ipsilateral SVZ wholemount (lower images; subep-ChAT^+^ neurons inhibition) vs. contralateral SVZ wholemount (upper images; (control)) from mice in panel E. Yellow arrows show EdU^+^/Ki67^+^ cells surrounding subep-ChAT^+^ neurons. Scale bar, 20 μm. **(H)** Upper; analysis of EdU^+^/GFAP^+^ cells per a subep-ChAT^+^ neurons in SVZ wholemounts of ipsilateral vs. contralateral in panel F. P < 0.039, t^15^ = 2.25, n=16, Paired t-test. Data collected from four stimulated *ChAT-Cre; AiD40* mice. Each dot represents total EdU^+^/GFAP^+^ cells surrounding a subep-ChAT^+^ neuron. Lower; analysis of EdU^+^/Ki67^+^ cells per a subep-ChAT^+^ neurons in SVZ wholemounts of ipsilateral vs. contralateral in panel G. P < 0.0164, t^15^ = 2.7, n=16, Paired t-test. Data collected from four stimulated *ChAT-Cre; AiD40* mice. Each dot represents total EdU^+^/Ki67^+^ cells surrounding a subep-ChAT^+^ neuron. **(I)** Experimental design (upper) and schematic representation (lower) of *in vivo* optogenetic activation for 6hrs post pAAV-hSyn-DIO-hM3D(Gq)-mCherry virus injection into ACC region of (P28) *CR-Cre* mice. **(J)** Immunofluorescence staining for ChAT (green), GFAP (gray) and EdU (purple) of ipsilateral SVZ wholemount (lower images; ACC-subep-ChAT^+^ circuit activation) vs. contralateral SVZ wholemount (upper images; (control)) from mice in panel I. Yellow arrows show EdU^+^/GFAP^+^ cells surrounding subep-ChAT^+^ neurons. Scale bar, 20 μm. **(K)** Immunofluorescence staining for ChAT (green), Ki67 (gray) and EdU (purple) of ipsilateral SVZ wholemount (lower images; ACC-subep-ChAT^+^ circuit activation) vs. contralateral SVZ wholemount (upper images; (control)) from mice in panel I. Yellow arrows show EdU^+^/Ki67^+^ cells surrounding subep-ChAT^+^ neurons. Scale bar, 20 μm. **(L)** Upper; analysis of EdU^+^/GFAP^+^ cells per a subep-ChAT^+^ neuron in SVZ wholemounts of ipsilateral vs. contralateral in panel B. P < 0.0046, t^15^ = 3.33, n=16, Paired t-test. Data collected from four stimulated *CR-Cre* mice. Each dot represents total EdU^+^/GFAP^+^ cells surrounding a subep-ChAT+ neuron. Lower; analysis of EdU^+^/Ki67^+^ cells per a subep-ChAT^+^ neuron in SVZ wholemounts of ipsilateral vs. contralateral in panel B. P < 0.0072, t^15^ = 3.11, n=16, Paired t-test. Data collected from four stimulated *CR-Cre* mice. Each dot represents total EdU^+^/Ki67^+^ cells surrounding a subep-ChAT+ neuron. All errors bars indicate SEM.

The activity of subep-ChAT^+^ neurons was evaluated by quantifying the levels of phospho-S6 ribosomal protein (P-S6), as depicted in **SI Appendix, Fig. S1A and C**. Stimulation of subep-ChAT^+^ neurons led to increased activity on the stimulated side compared to the control side. Conversely, inhibition of the subep-ChAT^+^ neurons resulted in the opposite effect, as illustrated in **SI Appendix, Fig. S1B and D**.

The activation and proliferation of LV NSCs were assessed by examining the colocalization of GFAP (quiescent NSC marker) and Ki67 (proliferation marker) with EdU. In the subep-ChAT^+^ activation experiments, the number of GFAP^+^/EdU^+^ cells surrounding subep-ChAT^+^ neurons in the stimulated (ipsilateral) SVZ were higher compared to the control (contralateral) SVZ **(Fig. 1 B and D; upper)**. Furthermore, the activated SVZ surrounding subep-ChAT^+^ neurons showed more Ki67^+^/EdU^+^ cells compared to the control SVZ **(Fig. 1 C and D; lower)**. In contrast, inhibition of subep-ChAT^+^ neurons resulted in a decrease in the number of GFAP^+^/EdU^+^ and Ki67^+^/EdU^+^ cells in the inhibited SVZ compared to their counterparts in the control SVZ **(Fig. 1 F, G and H)**.

We then used chemogenetics to activate the ACC-subep-ChAT^+^ circuit by injecting pAAV-hSyn-DIO-hM3D(Gq)-mCherry into the ACC of *CR-Cre* mice **(Fig. 1 I)**. The modulation of the cortical circuit lasted 6 hours, with EdU injection occurring 2 hours before the end of the experiment **(Fig. 1 I; upper and SI Appendix, Fig. S2 A; upper)**. Activation of this circuit resulted in an increased number of GFAP^+^/EdU^+^ cells adjacent to subep-ChAT^+^ neurons compared to the contralateral side **(Fig. 1 J and L; upper)**. Additionally, more Ki67^+^/EdU^+^ cells surrounding subep-ChAT^+^ neurons were observed compared to the contralateral side **(Fig. 1 J and L; lower)**. To inhibit this circuit, we used a Cre-dependent inhibitory opsin (AAV-hSyn1-SIO-stGtACR2-FusionRed) virus injected into the ACC of *CR-Cre* mice (**SI Appendix, Fig. S2 A)**. Conversely, inhibiting the ACC-subep-ChAT^+^ circuit led to a reduction in both GFAP^+^/EdU^+^ and/or Ki67^+^/EdU^+^ cells adjacent to subep-ChAT^+^ neurons on the inhibited side **(SI Appendix, Fig. S2 B, C and D)**. Upon activation and inhibition of the ACC-subep-ChAT^+^ circuit, the activity of subep-ChAT^+^ neurons exhibited an increase during circuit activation and a decrease during circuit inhibition, as illustrated in **SI Appendix, Fig. S3 A, B, C and D**. Taken together, these results strongly suggest that the ACC-subep-ChAT^+^ circuit can initiate the activation and proliferation of ventral LV qNSCs.

### The activation of cholinergic muscarinic 3 (M3) receptors initiate the activation and proliferation of LV qNSCs

To investigate the cholinergic regulatory pathway that initiates and facilitates the activation and proliferation of LV qNSCs, we initially evaluated the effects of carbachol (an ACh agonist) on the proliferative activity of LV qNSCs in a culture of SVZ NSCs. This was accomplished by inducing a quiescent state in LV NSCs first and subsequently applying carbachol for a 12-hour period during the proliferation stage. The levels of phosphorated-EGFR (pEGFR) in LV NSCs were quantified for carbachol-treated SVZ NSCs culture samples and compared to control culture samples **(Fig. 2 A)**. We observed an increase in the quantity of pEGFR protein in the carbachol-treated samples compared to the control samples **(Fig. 2 B)**. Following the same experimental design, we incubated both the treated and control samples with EdU for 15 minutes before the end of the experiment and subsequently stained them with a GFAP antibody **(Fig. 2 C)**. The number of GFAP^+^/EdU^+^ LV NSCs in the carbachol-treated samples was higher compared to the control samples **(Fig. 2 D)**. These results indicate that ACh plays a pivotal role in initiating the activation and facilitating the proliferation of LV qNSCs.

**Figure 2.**
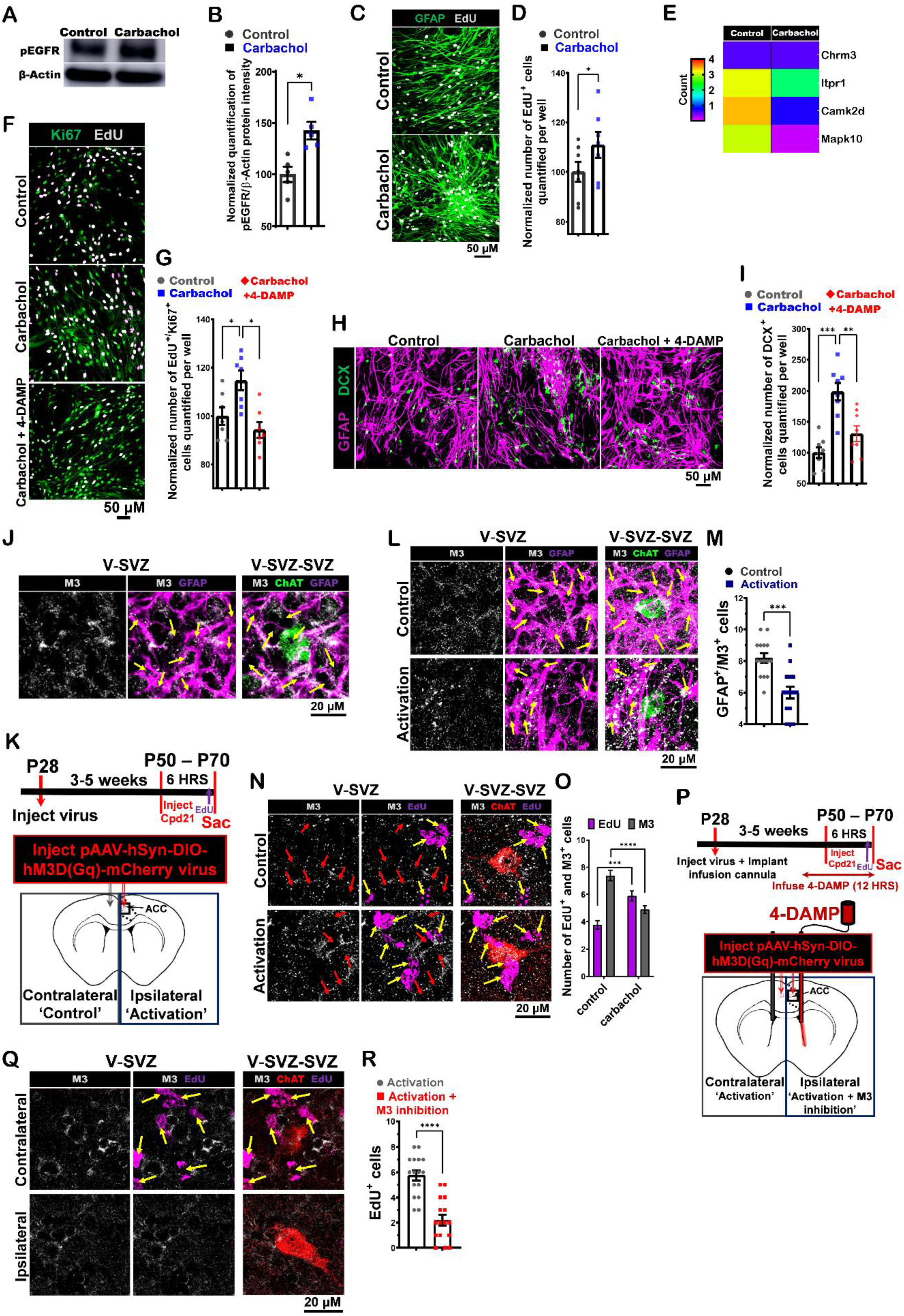
Modulation of activation and proliferation of LV qNSCs via M3 receptors. **(A)** Western blot detection of pEGFR and β-actin in control and treated samples with carbachol (15μM) of SVZ NSCs cultures in proliferation media collected after 12 hours. **(B)** Analysis normalized quantification of pEGFR/β-actin protein intensity for control and treated samples with carbachol from western plot in panel A. P = 0.0173, n=5, Paired t-test. **(C)** Immunofluorescence staining for GFAP (green) and EdU (gray) in control and treated samples with carbachol (15μM) of SVZ NSCs cultures in proliferation media collected after 12 hrs. Scale bar, 50 μm. **(D)** Analysis of normalized quantification of EdU^+^ cells per well in control and treated samples with carbachol (15μM) of SVZ NSCs cultures from panel C. P = 0.038, n=7, Paired t-test. **(E)** Heat map illustrating the RNA-Seq of differentially expressed transcripts from Chrm3, Itpr1, Camk2d and Mapk10 genes, induced by treating SVZ NSCs culture with carbachol (15μM) versus the control in proliferation media collected after 12 hrs. **(F)** Immunofluorescence staining for Ki67 (green) and EdU (gray) in treated samples with carbachol (15μM) and carbachol (15μM)/4-DAMP (3μM) versus control of SVZ NSCs cultures in proliferation media collected after 12 hrs. Scale bar, 50 μm. **(G)** Analysis of normalized quantification of EdU^+^/Ki67^+^ cells per well in treated samples with carbachol (15μM) and carbachol (15 μM)/4-DAMP (3 μM) versus control of SVZ NSCs cultures from panel F. P = 0.042 (Control vs. carbachol) and P = 0.021 (carbachol vs. carbachol (15μM)/4-DAMP), n=7, Paired t-test. **(H)** Immunofluorescence staining for GFAP (purple) and DCX (green) in treated samples with carbachol (15μM) and carbachol (15μM)/4-DAMP (3 μM) versus control of SVZ NSCs cultures in differentiation media collected after two days. **(I)** Analysis of normalized quantification of DCX^+^ cells per well in treated samples with carbachol (15μM) and carbachol (15μM)/4-DAMP (3 μM) versus control of SVZ NSCs cultures from panel H. P = 0.0004 (Control vs. carbachol) and P = 0.0094 (carbachol vs. carbachol/4-DAMP), n=7, Paired t-test. All errors bars indicate SEM. **(J)** Immunofluorescence staining for M3 (gray), ChAT (green) and GFAP (purple) in SVZ wholemount of C57BL/6J (P30) mice. Scale bar, 20 μm. **(K)** Experimental design (upper) and schematic representation (lower) of *in vivo* chemogenetic activation for 6 hours using pAAV-hSyn-DIO-hM3D(Gq)-mCherry virus injected into the ACC. **(L)** Immunofluorescence staining for M3 (gray), ChAT (green), and GFAP (purple) of ipsilateral side (lower images; ACC-subep-ChAT^+^ circuit activation) vs. contralateral side (upper images; (control)) from mice in panel C. Yellow arrows show GFAP^+^/M3^+^ cells surrounding a subep-ChAT^+^ neuron. Scale bar, 20 μm. **(M)** Analysis of GFAP^+^/M3^+^ cells per subep-ChAT^+^ neurons in SVZ wholemounts of ipsilateral vs. contralateral in panel D. P < 0.0009, t^14^ = 4.17, n=15, Paired t-test. Data collected from three stimulated *CR-Cre* mice. Each dot represents total GFAP^+^/M3^+^ cells surrounding a subep-ChAT^+^ neuron. **(N)** Immunofluorescence staining for M3 (gray), ChAT (green), and EdU (purple) of ipsilateral SVZ wholemount (lower images; ACC-subep-ChAT^+^ circuit activation) vs. contralateral SVZ wholemount (upper images; (control)) from mice in panel C. Yellow arrows show EdU^+^/M3^+^ cells surrounding a subep-ChAT^+^ neuron. Red arrows show EdU^+^/M3^+^ cells surrounding a subep-ChAT^+^ neuron. Scale bar, 20 μm. **(O)** Analysis of EdU^+^ and M3^+^ cells per subep-ChAT^+^ neuron in SVZ wholemounts of ipsilateral vs. contralateral in panel F. Two-way ANOVA (F (1,15) = 58.50; P < 0.0001), P = 0.0003 (EdU), P < 0.0001 (M3); n=16. Data collected from four stimulated *CR-Cre* mice. **(P)** Experimental design (upper) and schematic representation (lower) of *in vivo* chemogenetic activation for 6hrs post pAAV-hSyn-DIO-hM3D(Gq)-mCherry virus injection into ACC regions and cannulas implantation into LV regions of (P30) *CR-Cre* mice. **(Q)** Immunofluorescence staining for CHRM3 (gray), ChAT (red), and EdU (purple) of ipsilateral SVZ wholemount (lower images; ACC-subep-ChAT^+^ circuit activation and 4-DAMP infusion into ventral LV) vs. contralateral SVZ wholemount (upper images; ACC-subep-ChAT^+^ circuit activation (control)) from mice in panel H. Yellow arrows show EdU^+^ cells surrounding a subep-ChAT^+^ neuron. Scale bar, 20 μm. **(R)** Number of EdU^+^ cells per subep-ChAT^+^ neuron in SVZ wholemounts in panel H. P < 0.0001, t^15^ = 6.25, n=16, Paired t-test. Data collected from four stimulated *CR-Cre* mice. Each dot represents total EdU^+^ cells surrounding a subep-ChAT^+^ neuron.

To examine cholinergic receptor expression in LV qNSCs and explore potential differential gene expression associated with the molecular basis, we performed a bulk RNA sequencing (RNA-seq) experiment. Specifically, we compared carbachol-treated samples (12 hours) with control samples cultured in the proliferation media following the transition of LV NSCs into a quiescent state **(SI Appendix, Fig. S4 A)**. We observed a moderate level of M3 ACh receptor (CHRM3) gene expression in both samples, while the expression of other muscarinic receptors was minimal or absent **(Fig. S4 E)**. Although the expression of M3 receptors remained consistent between the two samples, our focus was to investigate their involvement in circuit activation due to their relatively high expression levels in the samples compared to other muscarinic receptors.

To understand the role of M3 receptors in the proliferation of LV NSCs, we treated SVZ NSCs cultures in proliferation media with either carbachol alone or carbachol in combination with 4-DAMP, a selective antagonist for M3. Subsequently, these cultures were then incubated with EdU and stained for Ki67 to detect proliferation **(Fig. 2 F)**. The number of EdU^+^/Ki67^+^ cells in both the control and carbachol (15 μM)/4-DAMP (3 μM) SVZ NSCs cultures was lower compared to the carbachol-treated culture **(Fig. 2 G)**.

In addition, SVZ NSCs cultures were treated with either carbachol alone or carbachol in combination with 4-DAMP for two days after a switch to differentiation media **(Fig. 2 H)**. The number of neuroblasts (DCX^+^ cells) was higher in the samples incubated with carbachol compared to both the control and carbachol/4-DAMP-incubated samples **(Fig. 2 I)**. Collectively, these findings suggest that the proliferative activation of LV NSCs can be regulated by the activity of the M3 receptors.

The expression of the CHRM3 gene was then confirmed by conducting quantitative reverse transcription PCR (RT-qPCR) analysis **(SI Appendix, Fig. S4 B)**. Both the RNAseq and qPCR datasets suggested that CHRM3 RNA is expressed in the LV qNSCs **(SI Appendix, Fig. S4 A and B)**. In addition to that, the co-staining of GFAP and M3 on the SVZ wholemount of *C57BL/6J* (P30) mice revealed the expression of M3 receptors in LV qNSCs surrounding subep-ChAT^+^ neurons **(Fig. 2 J)**.

To determine the role of M3 receptors in modulating ventral LV qNSCs through the ACC-subep-ChAT^+^ circuit, we activated this circuit using chemogenetic techniques, as illustrated in **Fig. 2 K**. pAAV-hSyn-DIO-hM3D(Gq)-mCherry virus was injected into the ACC region of *CR-Cre* mice. To identify proliferating cells, EdU was administered intraperitoneally 2 hours before the end of the experiment **(Fig. 2 K; upper)**. We observed a reduced number of GFAP^+^/M3^+^ cells surrounding subep-ChAT^+^ neurons on the activated side of the circuit when compared to the control side **(Fig. 2 L and M)**. Furthermore, as the number of EdU^+^ LV NSCs increased in the V-SVZ region due to circuit activation, there was a corresponding decrease in the level of M3 protein within the LV NSCs in the same region surrounding subep-ChAT^+^ neurons **(Fig. 2 N; lower and O)**. In contrast, on the control side, there were fewer EdU^+^ LV NSCs in the V-SVZ region surrounding the area of subep-ChAT^+^ cells **(Fig. 2 N; upper and O)**.

The activation of the cortical circuit has led to a decrease in M3 protein levels in the membrane of newly activated EdU^+^ LV NSCs **(Fig. 2 N)**. However, the RNA data did not indicate a decrease in the expression of CHRM3 in the samples treated with carbachol **(SI Appendix, Fig. S4 B)**. These findings suggest that the activation of the ACC-subep-ChAT^+^ circuit may result in the internalization of M3 receptors as the LV qNSCs are activated, and that reduced M3 receptor protein suggests removal of receptors from the cell surface (28).

To verify the importance of M3 receptors activation in modulating the activity of LV NSCs, we employed a selective inhibition strategy by locally infusing the M3 antagonist 4-DAMP into the ventral SVZ while simultaneously activating the ACC-subep-ChAT^+^ circuit **(Fig. 2 P)**. We found a significant decrease in the number of EdU^+^ LV NSCs adjacent to the subep-ChAT^+^ neurons on the inhibited side, in contrast to the side where the circuit was activated and multiple EdU^+^ LV NSCs were observed **(Fig. 2 Q and R)**. These findings provide further support for the critical role of M3 receptors in regulating the proliferative activity of LV qNSCs in the ventral SVZ.

### Control of the proliferation activity of ventral LV qNSCs through modulation of cortical circuitry via IP3R1 receptor signaling

As we examined the downstream signaling pathway associated with M3 receptors that regulate the proliferative activity of LV qNSCs, we initially analyzed RNAseq data from carbachol-treated and control samples **(SI Appendix, Fig. 2 A)**. During this analysis, we identified the ITPR1 (inositol 1,4,5-trisphosphate receptor type 1) gene, which exhibited higher expression in control samples but showed significant downregulation in carbachol-treated samples **(Fig. 2 E and SI Appendix, Fig. 2 C)**. The ITPR1 gene encodes the IP3R1 (inositol 1,4,5-trisphosphate receptor type 1) protein, known to function as second messengers in M3 activation (29). To confirm the presence of the IP3R1 protein in LV qNSCs, we examined its expression using immunofluorescence staining. This staining revealed that the IP3R1 protein colocalized with GFAP^+^ LV qNSCs, which surround subep-ChAT+ neurons in the ventral SVZ **(Fig. 3 A)**. These observations collectively indicate that the activation of ventral LV qNSCs may indeed be regulated through the activation of the IP3R1 receptor.

**Figure 3.**
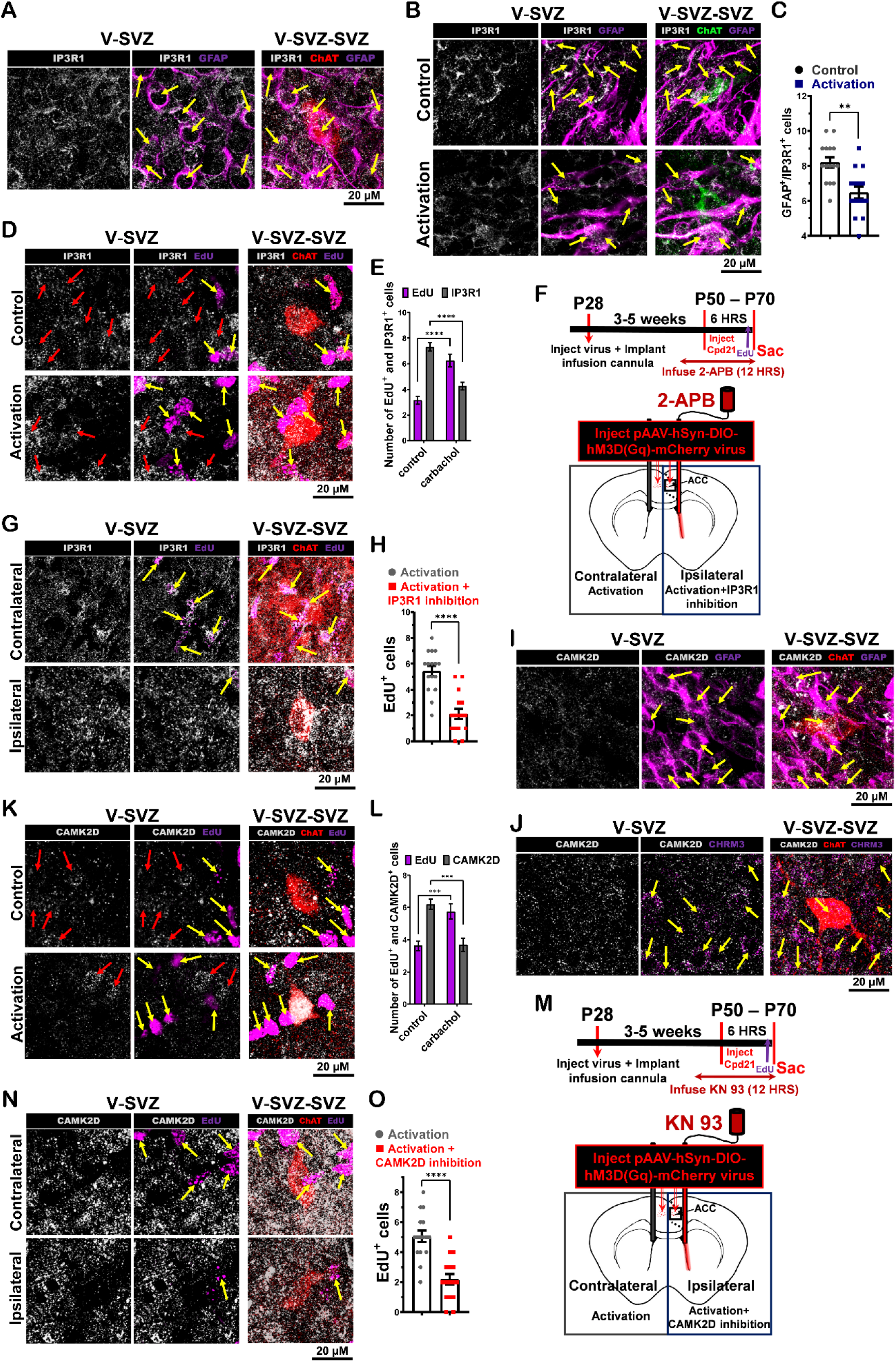
Control of proliferation activity of LV NSCs by IP3R1 and CAMK2D signaling *in vivo* following cortical circuit activation. **(A)** Immunofluorescence staining for IP3R1 (gray), ChAT (green) and GFAP (purple) in SVZ wholemount of C57BL/6J (P30) mice. Scale bar, 20 μm. **(B)** Immunofluorescence staining for IP3R1 (gray), ChAT (green), and GFAP (purple) of ipsilateral SVZ wholemount (lower images; ACC-subep-ChAT^+^ circuit activation) vs. contralateral SVZ wholemount (upper images; (control)) from mice in **Fig. 3C**. Yellow arrows show GFAP^+^/ IP3R1^+^ cells surrounding a subep-ChAT^+^ neuron. Scale bar, 20 μm. **(C)** Analysis of GFAP^+^/ IP3R1^+^ cells per a subep-ChAT^+^ neuron in SVZ wholemounts of ipsilateral vs. contralateral in panel C. P < 0.0010, t^14^ = 4.13, n=15, Paired t-test. Data collected from three stimulated *CR-Cre* mice. Each dot represents total GFAP^+^/ IP3R1^+^ cells surrounding a subep-ChAT^+^ neuron. **(D)** Immunofluorescence staining for IP3R1 (gray), ChAT (green), and EdU (purple) of ipsilateral SVZ wholemount (lower images; ACC-subep-ChAT^+^ circuit activation) vs. contralateral SVZ wholemount (upper images; (control)) from mice in **Fig. 3C**. Yellow arrows show EdU^+^/ IP3R1^+^ cells surrounding a subep-ChAT^+^ neuron. Red arrows show EdU^-^/ IP3R1^+^ cells surrounding a subep-ChAT^+^ neuron. Scale bar, 20 μm. **(E)** Analysis of EdU^+^ and IP3R1^+^ cells per subep-ChAT^+^ neuron in SVZ wholemounts of ipsilateral vs. contralateral in panel E. Two-way ANOVA (F (1,15) = 76.29; P < 0.0001), P < 0.0001 (EdU), P < 0.0001 (IP3R1); n=16. Data collected from four stimulated *CR-Cre* mice. **(F)** Experimental design (upper) and schematic representation (lower) of *in vivo* chemogenetic activation for 6hrs post pAAV-hSyn-DIO-hM3D(Gq)-mCherry virus injection into ACC regions and cannulas implantation into LV regions of (P30) *CR-Cre* mice. **(G)** Immunofluorescence staining for IP3R1 (gray), ChAT (red), and EdU (purple) of ipsilateral SVZ wholemount (lower images; ACC-subep-ChAT^+^ circuit activation and 2-APB infusion into ventral LV) vs. contralateral SVZ wholemount (upper images; ACC-subep-ChAT^+^ circuit activation (control)) from mice in panel G. Yellow arrows show EdU^+^ cells surrounding a subep-ChAT^+^ neuron. Scale bar, 20 μm. **(H)** Number of EdU^+^ cells per subep-ChAT^+^ neuron in SVZ wholemounts in panel G. Paired t-test p < 0.0001, n=16, from 4 stimulated *CR-Cre* mice. Each dot represents the number of EdU^+^ cells surrounding a subep-ChAT^+^ neuron. All errors bars indicate SEM. **(I)** Immunofluorescence staining for CAMK2D (gray), ChAT (red) and GFAP (purple) in SVZ wholemount of C57BL/6J (P30) mice. Scale bar, 20 μm. **(J)** Immunofluorescence staining for CAMK2D (gray), ChAT (red) and CHRM3 (purple) in SVZ wholemount of C57BL/6J (P30) mice. Scale bar, 20 μm. **(K)** Immunofluorescence staining for CAMK2D (gray), ChAT (red), and EdU (purple) of ipsilateral SVZ wholemount (lower images; ACC-subep-ChAT^+^ circuit activation) vs. contralateral SVZ wholemount (upper images; (control)) from mice in **Fig. 3C**. Yellow arrows show EdU^+^/ CAMK2D^+^ cells surrounding a subep-ChAT^+^ neuron. Red arrows show EdU^-^/ CAMK2D^+^ cells surrounding a subep-ChAT^+^ neuron. Scale bar, 20 μm. **(L)** Analysis of EdU^+^ and CAMK2D^+^ cells per subep-ChAT^+^ neuron in SVZ wholemounts of ipsilateral vs. contralateral in panel D. Two-way ANOVA (F (1,15) = 50.45; P < 0.0001), P = 0.0007 (EdU), P = 0.0001 (CAMK2D); n=16. Data collected from four stimulated *CR-Cre* mice. **(M)** Experimental design (upper) and schematic representation (lower) of *in vivo* chemogenetic activation for 6hrs post pAAV-hSyn-DIO-hM3D(Gq)-mCherry virus injection into ACC regions and cannulas implantation into LV regions of (P30) *CR-Cre* mice. **(N)** Immunofluorescence staining for CAMK2D (gray), ChAT (red), and EdU (purple) of ipsilateral SVZ wholemount (lower images; ACC-subep-ChAT^+^ circuit activation and KN 93 infusion into ventral LV) vs. contralateral SVZ wholemount (upper images; ACC-subep-ChAT^+^ circuit activation (control)) from mice in panel F. Yellow arrows show EdU^+^ cells surrounding a subep-ChAT^+^ neuron. Scale bar, 20 μm. **(O)** Number of EdU^+^ cells per subep-ChAT^+^ neuron in SVZ wholemounts in panel G. P < 0.0001, t^15^ = 5.668, n=16, Paired t-test. Data collected from four stimulated *CR-Cre* mice. Each dot represents total EdU^+^ cells surrounding a subep-ChAT^+^ neuron.

To examine the role of IP3R1 receptors, we investigated how modulation of the ACC-subep-ChAT^+^ circuit affects their activity in LV NSCs. When the cortical circuit was stimulated, we observed a decrease in the number of GFAP^+^/IP3R1^+^ cells in the ventral SVZ surrounding subep-ChAT^+^ neurons on the activated side compared to the control side **(Fig. 3 B and C)**. Furthermore, while the number of EdU^+^ LV NSCs surrounding subep-ChAT^+^ neurons increased on the activated side of the circuit, the number of IP3R1^+^ LV qNSCs decreased **(Fig. 3 D; lower and E)**. In contrast, the control ventral SVZ showed the opposite pattern, indicating that most LV NSCs were in a quiescent state and characterized by the expression of the IP3R1 protein **(Fig. 3 D; upper and E)**. However, the activated LV qNSCs were identified as EdU^+^/IP3R1^-^cells, suggesting that the ACC-subep-ChAT^+^ circuit controls the proliferative activity of ventral LV NSCs by upregulating IP3R1 signaling.

Furthermore, we examined the role of IP3R1 receptors in LV NSCs modulation by concurrently activating the circuit and selectively inhibiting IP3R1 receptors using the selective antagonist 2-APB **(Fig. 3 F)**. With circuit activation and IP3R1 receptor inhibition, we observed that the number of EdU^+^ LV NSCs was rarely seen around subep-ChAT^+^ neurons in the ventral SVZ. In contrast, with circuit activation alone several EdU^+^ LV NSCs were observed **(Fig. 3 G and H)**. Altogether, these data suggest that IP3R1 receptor signaling is necessary for ACC-subep-ChAT^+^ circuit activation of ventral LV NSCs surrounding subep-ChAT^+^ neurons.

### Regulation of the proliferative activity of ventral LV qNSCs through modulation of cortical circuitry via the CAMK2D signaling pathway

The activation of the cortical circuit involving CHRM3 and ITPR1 receptors in LV qNSCs can induce the release of intracellular calcium from the endoplasmic reticulum (ER) (30). This increase in cytoplasmic calcium levels can lead to the activation of CaM Kinase signaling (31). We then investigated the differential gene regulation of this signaling in our RNAseq data, which revealed a significant decrease in CAMK2D gene expression in carbachol-treated samples compared to the control group. This finding was further confirmed by RT-qPCR experiments **(Fig. 2 E and SI Appendix, Fig. S4 D)**. Immunostaining in the V-SVZ demonstrated the expression of CAMK2D protein in GFAP^+^ LV qNSCs surrounding subep-ChAT^+^ neurons **(Fig. 3 I)**. Additionally, the CAMK2D protein was observed to colocalize with M3 receptors on LV qNSCs within the ventral SVZ **(Fig. 3 J)**. These findings collectively suggest that CAMK2D signaling acts downstream of ACC-subep-ChAT^+^ circuit modulation in LV qNSCs adjacent to subep-ChAT^+^ neurons in the ventral SVZ. To test this possibility, we examined the modulation of cortical circuit in ventral LV qNSCs and its relationship with the signaling activity of CAMK2D protein. After 6 hours of circuit activation, we observed a lower number of GFAP^+^/CAMK2D^+^ LV NSCs adjacent to subep-ChAT^+^ neurons in the activated SVZ compared to their counterparts in the control SVZ **(SI Appendix, Fig. S5A and B)**. Additionally, there was an increase in the number of EdU^+^ LV NSCs accompanied by a simultaneous decrease in the number of CAMK2D^+^ LV NSCs surrounding subep-ChAT^+^ neurons in the activated SVZ **(Fig. 3 K and L)**. These findings suggest that activating the cortical circuit may regulate the proliferation activity of ventral LV NSCs through CAMK2D signaling.

We simultaneously activated the circuit and locally inhibited the CAMK2D protein using the selective inhibitor KN 93. Subsequently, we assessed the impact on the proliferation of LV NSCs surrounding subep-ChAT^+^ neurons **(Fig. 3 M)**. The results showed a reduction in the number of EdU^+^ cells neighboring subep-ChAT^+^ neurons on the ipsilateral side (i.e., circuit activation with local CAMK2D inhibition) compared to their counterparts on the contralateral side (i.e., circuit activation only) **(Fig. 3 N and O)**. These findings strongly support the role of the CAMK2D signaling pathway in the mechanism by which the ACC-subep-ChAT^+^ circuit regulates the proliferation of LV NSCs in the ventral SVZ.

### Regulation of the proliferative activity of ventral LV NSCs through modulation of cortical circuitry via the MAPK10 signaling pathway

Thus, activation of the ACC-subep-ChAT^+^ circuit upregulates CAMK2D signaling in ventral LV NSCs, potentially activating downstream proliferation signaling pathways (32). Screening the RNAseq dataset of carbachol-treated samples compared to the control revealed a significant downregulation of the MAPK10 gene **(SI Appendix, Fig. S4 A and Fig. 2 E)**. This suggests the involvement of the MAPK10 pathway in modulating the proliferation of ventral LV NSCs through the cortical circuit. Cellular expression analysis showed the presence of MAPK10 protein in both GFAP^+^ LV NSCs and CHRM3^+^ cells in the ventral V-SVZ, neighboring subep-ChAT^+^ neurons **(Fig. 4 A and B)**. These observations suggest that the ACC-subep-ChAT^+^ circuit plays a role in regulating the proliferation process of ventral LV NSCs by activating the MAPK10 signaling pathway.

**Figure 4.**
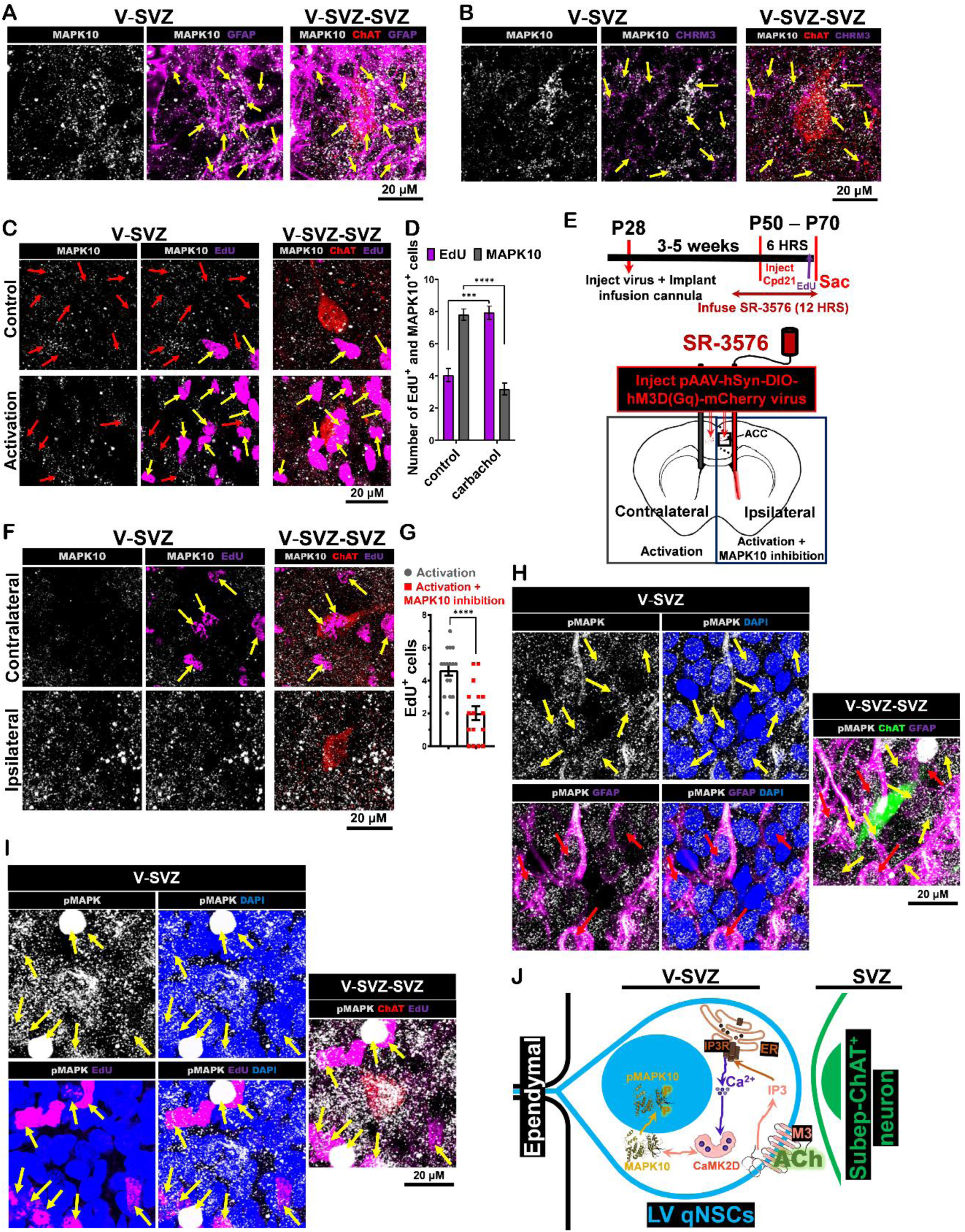
*In vivo* control of proliferation of LV qNSCs by MAPK10 signaling following cortical circuit activation. **(A)** Immunofluorescence staining for MAPK10 (gray), ChAT (red) and GFAP (purple) in SVZ wholemount of C57BL/6J (P30) mice. Scale bar, 20 μm. **(B)** Immunofluorescence staining for MAPK10 (gray), ChAT (red) and CHRM3 (purple) in SVZ wholemount of C57BL/6J (P30) mice. Scale bar, 20 μm. **(C)** Immunofluorescence staining for MAPK10 (gray), ChAT (red), and EdU (purple) of ipsilateral SVZ wholemount (lower images; ACC-subep-ChAT^+^ circuit activation) vs. contralateral SVZ wholemount (upper images; (control)) from mice in **Fig. 3C**. Yellow arrows show EdU^+^/ MAPK10^+^ cells surrounding a subep-ChAT^+^ neuron. Red arrows show EdU^-^/ MAPK10^+^ cells surrounding a subep-ChAT^+^ neuron. Scale bar, 20 μm. **(D)** Analysis of EdU^+^ and MAPK10^+^ cells per subep-ChAT^+^ neuron in SVZ wholemounts of ipsilateral vs. contralateral in panel D. Two-way ANOVA (F (1,15) = 54.19; P < 0.0001), P = 0.0005 (EdU), P < 0.0001 (MAPK10); n=16. Data collected from four stimulated *CR-Cre* mice. **(E)** Experimental design (upper) and schematic representation (lower) of *in vivo* chemogenetic activation for 6hrs post pAAV-hSyn-DIO-hM3D(Gq)-mCherry virus injection into ACC regions and cannulas implantation into LV regions of (P30) *CR-Cre* mice. **(F)** Immunofluorescence staining for MAPK10 (gray), ChAT (red), and EdU (purple) of ipsilateral SVZ wholemount (lower images; ACC-subep-ChAT^+^ circuit activation and SR-3576 infusion into ventral LV) vs. contralateral SVZ wholemount (upper images; ACC-subep-ChAT^+^ circuit activation (control)) from mice in panel F. Yellow arrows show EdU^+^ cells surrounding a subep-ChAT^+^ neuron. Scale bar, 20 μm. **(G)** Number of EdU^+^ cells per subep-ChAT^+^ neuron in SVZ wholemounts in panel G. P < 0.0001, t^15^ = 5.65, n=16, Paired t-test. Data collected from four stimulated *CR-Cre* mice. Each dot represents total EdU^+^ cells surrounding a subep-ChAT^+^ neuron. All errors bars indicate SEM. **(H)** Immunofluorescence staining for pMAPK (gray), ChAT (green), EdU (purple) and DAPI (blue) in the SVZ wholemount of the ACC-subep-ChAT^+^ circuit activation from mice in **Fig. 3C**. Yellow arrows show GFAP^-^/pMAPK^+^ cells surrounding a subep-ChAT^+^ neuron. Red arrows show GFAP^+^ cells surrounding a subep-ChAT+ neuron. Scale bar, 20 μm. **(I)** Immunofluorescence staining for pMAPK (gray), ChAT (red), EdU (purple) and DAPI (blue) in the SVZ wholemount of the ACC-subep-ChAT^+^ circuit activation from mice in **Fig. 3C**. Yellow arrows show EdU^+^/pMAPK^+^ cells surrounding a subep-ChAT^+^ neuron. Scale bar, 20 μm. **(J)** Illustration depicting the molecular mechanism through which the ACC-subep-ChAT^+^ circuit regulates the proliferation of LV qNSCs.

We examined the impact of cortical circuit activation on MAPK10 signaling. The results revealed a decrease in the number of GFAP^+^/MAPK10^+^ cells on the activated side compared to the control side **(SI Appendix, Fig. S6A and B)**. Additionally, there was an increase in the number of EdU^+^ LV NSCs and a decrease in the number of MAPK10^+^ cells on the activated side compared to the control side **(Fig. 4 C and D)**.

Furthermore, we examined the necessity of MAPK10 signaling in regulating the proliferation of LV qNSCs within the ventral SVZ in response to cortical circuit activation. This was accomplished by administering SR-3576, a selective MAPK10 inhibitor, within the ventral V-SVZ simultaneously with the activation of the cortical circuit **(Fig. 4 E)**. Notably, we observed a significant reduction in the number of EdU^+^ LV NSCs in the ipsilateral SVZ (i.e., circuit activation with local MAPK10 inhibition) compared to the relatively higher number of EdU^+^ LV NSCs in the contralateral SVZ (i.e., circuit activation alone) **(Fig. 4 F and G)**.

We proceeded to examine the phosphorylation-induced activation of MAPK10 and its subsequent nuclear localization within LV NSCs following the activation of the cortical circuit. The results have shown that several GFAP^+^ LV qNSCs initiate proliferation in the V-SVZ area surrounding subep-ChAT^+^ neurons. These cells exhibit phosphorylated MAPK (pMAPK) that relocates into the nucleus after cortical circuit activation **(Fig. 4 H)**. Furthermore, this has been confirmed by demonstrating the colocalization of EdU with pMAPK within the nuclei of the activated LV qNSCs in the V-SVZ region surrounding subep-ChAT^+^ neurons following ACC-subep-ChAT^+^ activation **(Fig. 4 I)**. These results indicate that the proliferation activity of LV NSCs in the ventral V-SVZ, particularly around subep-ChAT^+^ neurons, is regulated through MAPK signaling.

Taken together these findings confirm the essential role of the MAPK10 signaling pathway in mediating the modulation of ventral LV qNSCs adjacent to subep-ChAT^+^ neurons through the ACC-subep-ChAT^+^ circuit.

## Discussion

The study presented here investigates the molecular basis of a cortical circuit involved in the regulation of activation and proliferation of LV qNSCs in the SVZ. Our findings demonstrate that modulation of the ACC-subep-ChAT^+^ circuit for a brief period of 6 hours can exert control over LV qNSCs activation and proliferation, which is closely associated with subep-ChAT^+^ neurons in the ventral SVZ. The subep-ChAT^+^ neurons, a cholinergic population in the SVZ, regulate LV NSCs proliferation in an activity-dependent manner (26). Furthermore, these local cholinergic neurons receive inputs from both local GABAergic and cingulate glutamatergic neurons, which play a role in regulating SVZ neurogenesis in the ventral SVZ (27). Consequently, this study sheds light on the role of cholinergic signaling and the specific molecular mechanisms involved in the regulation of ventral LV NSCs activation and proliferation.

The incubation of postnatal SVZ NSC cultures with carbachol, an ACh agonist, enhanced the early-stage proliferation of LV qNSCs. The M3 muscarinic receptors were critical in regulating LV qNSCs near subep-ChAT^+^ neurons in the ventral SVZ. However, the impact of M3 modulation on non-excitable cells like LV qNSCs has been poorly studied and requires further research.

Additionally, the activation of the ACC-subep-ChAT^+^ circuit was found to reduce the protein levels of M3 receptors in newly proliferative (EdU^+^) ventral LV NSCs **(Fig. 3 C)**, suggesting receptor internalization as LV qNSCs become activated (28). Future research could contribute to a deeper understanding of the mechanisms behind M3 receptors internalization, phosphorylation, and desensitization in ventral LV qNSCs following ACC-subep-ChAT^+^ circuit modulation.

The cholinergic system involving the M3 receptor has also been shown to play a significant role in hippocampal neurogenesis and is implicated in the cognitive deficits observed in Alzheimer’s disease, as well as hippocampus-dependent learning and memory (33). Studies have demonstrated that the M3 receptor is activated in the hippocampus following agonist treatment and fear conditioning (34). The M3 receptors may potentially play a critical role in SVZ-olfactory neurogenesis through olfactory perceptual learning behavior. Further research could be directed towards providing valuable insights into studying the contributions of the M3 signaling pathway-mediated SVZ-olfactory neurogenesis in behavior associated with olfactory learning.

The M3 receptor activates various signaling pathways, including the formation of inositol 1,4,5-trisphosphate (IP3) through phospholipase C (PLC). IP3 then binds to IP3 receptors (IP3Rs), triggering the release of Ca^2+^ from the endoplasmic reticulum (ER) (35, 36). Our study suggests a downregulation of the ITPR1 gene expression (encoding IP3R), based on RNAseq and RT-qPCR data. While the role of IP3R in stem cell proliferation has not been extensively studied, it has been shown to be implicated in apoptosis in mouse embryonic stem cells (37). This suggests that IP3Rs may contribute to the early stages of stem cell proliferation. However, the specific roles of IP3Rs in the SVZ remain obscure (38).

The study’s other significant findings include a decrease in expressions of CAMK2D and MAPK10 following carbachol treatment in postnatal SVZ NSC cultures. CAMK2D, a multifunctional Ca^2+^/calmodulin (CaM)-dependent protein kinase regulated by intracellular Ca^2+^ signaling (39), plays a critical role in the proliferation of LV qNSCs situated adjacent to subep-ChAT^+^ neurons **(Fig. 4)**. Previous research has primarily focused on CAMK2D and postnatal/adult neurogenesis in the hippocampus, where its involvement in neural plasticity, memory processing, and learning has been demonstrated (40).

The present study contributes to the understanding of CAMK2D’s involvement in LV qNSCs proliferation by implicating the MAPK signaling pathway, in accord with previous findings (41). Earlier investigations showed that CAMK2 can activate MAPK signaling (42), while MAPKs can regulate CAMK2 proteins activity (43). Despite the limited research on MAPK and neural stem cells (44), a study has presented evidence of MAPK10’s role in neuronal differentiation in human neural stem cells derived from both ESCs and iPSCs (45). Taken together, these findings collectively provide support for the functional significance of CAMK2D and MAPK10 in regulating ventral LV NSC proliferation, emphasizing its relationship with the ACC-subep-ChAT^+^ circuit.

The activation of MAPK is initiated through phosphorylation, and its subsequent translocation into the nucleus has previously been linked to cellular proliferation (46). Our present study highlights the crucial role of the MAPK10 protein’s translocation into the nucleus of LV qNSCs, functioning as a controller of proliferation activity. This regulation is triggered by the activation of the ACC-subep-ChAT+ circuit.

The study further revealed the expression of M3, IP3R1, CAMK2D, and MAPK10 proteins in GFAP^+^ LV qNSCs located adjacent to subep-ChAT^+^ neurons **(Fig. 4 J)**. By conducting *in vivo* circuit modulation and utilizing targeted protein inhibitors, we determined their contributions to the activation and proliferation of LV qNSCs. These findings advance our understanding of the molecular mechanisms governing the regulation of ventral LV qNSCs by cortical circuits.

In humans, active neurogenesis has been observed at the wall of the lateral ventricles. This process generates migratory neuroblasts during the very early years after birth only (47). However, the molecular mechanisms regulating this circuitry in humans have not been studied yet. By examining the analogous process in rodents, our study paves the way for new research into the significance of signaling pathways associated with cortical activity-dependent neurogenesis in the SVZ. This research pertains to both postnatal and adult animals under normal physiological conditions, including possible associated disorders. Additionally, it may offer insights for future studies into the neural circuitry that regulates early human postnatal neurogenesis in the SVZ niche, along with its molecular mechanisms of functional contribution during brain development.

## Materials and Methods

### Material

#### Chemicals

The antibodies used in this paper are as follows: BMP-4 Protein (Catalog #: 5020-BP) R&D Systems, Carbachol (Catalog # 212385-M) Millipore Sigma, Compound 21 (DREADD Agonist 21 dihydrochloride) (Catalog # SML2392) Sigma-Aldrich, 4-DAMP (Catalog # SML0255) Sigma-Aldrich, 2-APB (CAS 524-95-8) Calbiochem, KN-93 (ab120980) abcam, SR-3576 (CAS 1164153-22-3) Calbiochem.

#### Mice

All mouse experiments were performed according to an approved protocol by the Institutional Animal Care and Use Committee at Duke University. The mice were group-housed on a standard 12-hour light/dark cycle, with lights on at 7 a.m. The housing maintained a controlled average ambient temperature of 21°C and 45% humidity. The following mouse lines were purchased from JAX: *C57BL/6J* (000664); *ChAT-ChR2-YFP* (014545); *ChAT-Cre* (006410); *Ai40D* (021188) and *Cr-Cre* (010774).

#### Viruses

pAAV-hSyn-DIO-hM3D(Gq)-mCherry (addgene #44361).

pAAV_hSyn1-SIO-stGtACR2-FusionRed (addgene #105677).

#### Antibodies

The antibodies used in this paper are as follows: GFP (Catalog # GFP-1020) AVES, Anti-Choline Acetyltransferase Antibody (Catalog # AB144P) Millipore sigma, Anti-tdTomato [16D7] Antibody (EST203). Kerafast, INC, RFP Antibody (Catalog # 600-401-379) Rockland, RFP antibody (Catalog # ab62341) Abcam, Ki67 (Catalog # ab15580) Abcam, Ki67 (Catalog # CPCA-Ki67) EnCor Biotechnology, Ki-67 Antibody (SolA15) (Catalog # 14-5698-82) ThermoFisher Scientific, beta Actin antibody (BA3R), HRP (Catalog # MA515739HRP) ThermoFisher Scientific, HRP-conjugated Beta Actin antibody (Catalog # HRP-60008) Proteintech, F(ab’)_2_ Fragment Goat Anti-Guinea Pig IgG (Catalog # 106-006-003) Jackson ImmunoResearch, F(ab’)_2_ Fragment Goat Anti-Rabbit IgG (Catalog # 111-036-003) Jackson ImmunoResearch, GFAP Chicken antibody (Catalog # GFAP) Aves Labs, CHRM3 antibody (Catalog # PA5-85322) ThermoFisher Scientific, CHRM3 antibody (Catalog # MAB6378SP) R&D Systems, CHRM3 antibody (Catalog # 580011) Novus Biologicals, ITPR1 antibody (Catalog # PA5-85753) ThermoFisher Scientific, CAMK2D antibody (Catalog # 15443-1-AP) Thermo Fisher Scientific, CAMK2D antibody (Catalog # H00000817-M02) Abnova, MAPK10 Polyclonal Antibody (Catalog # 17572-1-AP) ThermoFisher Scientific. Phospho-JNK1/JNK2/JNK3 (Thr183, Thr221) (Catalog # MA5-32159) ThermoFisher Scientific.

### Experiment

#### Stereotaxic injections

Stereotaxic injections were performed while mice were kept deeply anesthetized in a stereotaxic frame (David Kopf Instruments) using isoflurane. To modulate the activity of subep-ChAT^+^ neuron, optical fibers were implanted into the LVs within the ventral SVZ of *ChAT-ChR2-EYFP* (P30) and *ChAT-Cre; AiD40* (P30) mice as the following coordinates relative to Bregma: anteroposterior (AP): +0.8, mediolateral (ML): ±0.65, dorsoventral (DV): 2.1 from the brain surface. For *in vivo* chemogenetics activation of the ACC-subep-ChAT^+^ circuit, pAAV-hSyn-DIO-hM3D(Gq)-mCherry (300nl) virus was injected into the ACC region of *Cr-Cre* (P30) mice at the following coordinates relative to Bregma: AP: +0.8, ML ± 0.25, DV: 0.3 from the brain surface. To achieve *in vivo* optogenetic inhibition of the ACC-subep-ChAT^+^ circuit, AAV-hSyn1-SIO-stGtACR2-FusionRed virus (300nl) was injected into the ACC region of *Cr-Cre* (P30) mice as the following coordinates relative to Bregma: AP: +0.8, ML ± 0.25, DV: 0.3 from brain surface. All viruses were infused slowly for over 10 minutes using Nanoject (Drummond Scientific) connected to a glass pipette. The injection pipette was left in place for 10 minutes post-injection before being retracted.

For the pharmacological testing of ventral subventricular zone (SVZ) local inhibition of CHRM3, IPTR1, CAMK2D, and MAPK10 using 4-DAMP, 2-APB, KN-93, and SR-3576, respectively, we utilized a micro-osmotic pump (Model 1003D, alzet). A cannula was implanted into LV of *Cr-Cre* (P30) mice at the coordinates AP: +0.8, ML ± 0.65, DV: 2.1 from brain surface. This procedure was performed simultaneously with the injection of pAAV-hSyn-DIO-hM3D(Gq)-mCherry (300nl) virus into ACC as described in the above paragraph. The cortical circuit was activated by intraperitoneal injection of cpd21 6 hours after connecting the osmotic pump, which allowed for drug perfusion. The manufacturer’s instructions were followed, with the drug being perfused at a rate of 1 μl per hour for a duration of 12 hours, as indicated in **Figs. 3 and 4**.

#### Immunofluorescence staining and imaging

Brain tissues for IHC staining were prepared for subep-ChAT^+^ neurons and cortical circuit modulations in *in vivo* optogenetic and *in vivo* chemogenetics experiments. At the endpoint of the experiments, mice were deeply anesthetized with isoflurane and perfused transcardially with phosphate-buffered saline (PBS), followed by 4% paraformaldehyde (PFA) in PBS. The perfused brains were then removed and postfixed overnight at 4°C in 4% PFA.

The SVZ wholemounts were dissected out from the brains and incubated in a blocking solution containing 5% donkey serum and TBST for 100 minutes at room temperature. They were then incubated overnight at room temperature in PBS containing 1% donkey serum and specific antibodies, which had been validated in our previous publication (27) or by publications available on the vendor’s website for each antibody. After incubation, the SVZ wholemounts were washed with PBS and incubated with secondary antibodies, such as Alexa-594 (1:1000, LifeTech), Alexa-488 (1:1000, LifeTech), or Alexa-647 (1:1000, LifeTech), for 2 hours at room temperature, followed by washing with PBS. Sections were counterstained with a 4’,6-diamidino-2-phenylindole solution (DAPI; Sigma-Aldrich). After four washes with TBST, the sections were coverslipped with Fluoromount (Sigma), an aqueous mounting medium.

Images were captured using a Leica SP8 upright confocal microscope (Zeiss) with 10 ×, 20 ×, and 40 × objectives through tile scan imaging, controlled by Zen software (Zeiss).

#### *In vivo* optogenetic circuit modulation

To target the LVs of *ChAT-ChR2-EYFP* (P30) and *ChAT-Cre; AiD40* (P30) mice at the coordinates AP: +0.8, ML: ±0.65, DV: 2.1 from the brain surface, optical fibers were placed under isoflurane anesthesia. This was achieved using implantable mono fiber-optic fibers (200 μm, 0.22 NA, Doric). The protruding ferrule end of the cannula was then connected to a fiber cord, which in turn was attached to a rotary coupling joint (Doric) to allow free movement for the animals. Three to five weeks after viral infection, light stimulation for circuit activation was delivered using TTL control (Master 8, AMPI) of a 473-nm laser (IkeCool). The stimulation consisted of 10 ms pulses at 10 Hz, lasting 10 seconds, and was given once every minute. Additionally, for circuit inhibition, light stimulation was delivered using TTL control (Master 8, AMPI) of a 556-nm laser (IkeCool). In this case, a continuous laser beam with a power of 2 mW was used to avoid overheating in the local area.

In addition, mono fiber-optic fibers (200 μm, 0.22 NA, Doric) were implanted under isoflurane anesthesia to target the LV regions for the purpose of targeting the ventral SVZ at the coordinates AP: +0.9, ML: ±0.65, DV: 2.1 from the brain surface. This procedure was performed after the injection of the AAV-hSyn1-SIO-stGtACR2-FusionRed virus in P30 *CR-Cre* mice, as described in **SI Appendix, Fig. S2A**. Following a period of three to five weeks after viral infection, continuous light stimulation was delivered using TTL control (Master 8, AMPI) of a 473-nm laser (IkeCool). Only the LV region on the ipsilateral side was stimulated, utilizing a laser power of 2 mW.

To analyze ChAT, Ki67, GFAP, EdU, CHRM3, ITPR1, CAMK2D, and MAPK10, we dissected SVZ wholemounts and specifically chose areas surrounding the activated or inhibited subep-ChAT^+^ neurons for further analysis.

#### *In vivo* stimulation of chemogenetics circuits and inhibition of local proteins in the SVZ

After performing stereotaxic viral infusion as described in the “Stereotaxic Injections” section and the experimental design outlined in **Fig. 1 I**, Compound 21 (Cpd21; DREADD Agonist 21 dihydrochloride (SML2392)) from Millipore Sigma was dissolved in 0.9% saline and stored at - 20°C until it was used, as shown in **Fig. 1, 3, and 4**. Intraperitoneal injections of Cpd21 were administered at a volume of 1 mg/kg. The *in vivo* chemogenetic circuit stimulation was initiated by injecting Cpd21 intraperitoneally for 6 hours prior to the experiment’s conclusion. Additionally, simultaneous testing of *in vivo* chemogenetic circuit stimulation and local inhibition of CHRM3, ITPR1, CAMK2D, and MAPK10 proteins in the lateral ventricles (LVs) was performed, as depicted in **Figs. (3 H and Q) and (4 F, and N)**. In these experiments, the ventral SVZ was subjected to local protein inhibition using 4-DAMP (30 nM), 2-APB (60 μM), KN 93 (0.5 μM), and SR-3576 (25 nM), respectively, for 12 hours. The circuit was then activated in the last 6 hours before the experiment concluded.

For the analysis of EdU, GFAP, Ki67, CHRM3, ITPR1, CAMK2D, and MAPK10, SVZ wholemounts were dissected, and the number of EdU^+^/GFAP^+^, CHRM3^+^/GFAP^+^, ITPR1^+^/GFAP^+^, CAMK2D^+^/GFAP^+^, MAPK10^+^/GFAP^+^, EdU^+^/Ki67^+^, CHRM3^+^/Ki67^+^, ITPR1^+^/Ki67^+^, CAMK2D^+^/Ki67^+^, and MAPK10^+^/Ki67^+^ cells surrounding subep-ChAT^+^ neurons on the ipsilateral (circuit activation or circuit activation/local proteins inhibition) sides were compared to the contralateral sides (control).

#### *In vivo* EdU staining

The staining of 5-ethynyl-2’-deoxyuridine (EdU) was performed using the EdU Cell Proliferation Kit for Imaging (EdU *in vivo* Kits) from baseclick GmbH, Germany, following the manufacturer’s protocol. EdU was prepared at a concentration of 50 mg/20 mL in sterile PBS and used for pulse labeling of adult mice by administering an intraperitoneal (IP) injection of 500 μL of the dissolved EdU (50 mg/kg). The mice were then harvested according to the procedures described in Figures 1, 3, 4, 5, and 6.

For EdU labeling of the intended SVZ wholemounts, a sequential staining approach was employed. First, the wholemounts were stained for other antibodies as described in the immunofluorescence staining section. Subsequently, they were counterstained with DAPI. The sections were then washed three times with 3% BSA in PBS and incubated for 30 minutes in a reaction cocktail consisting of deionized water, reaction buffer, catalyst solution, dye azide, and buffer additive. During this incubation, the sections were protected from light. After the reaction cocktail was removed, the wholemounts were washed three times with 3% BSA in PBS. Finally, they were mounted in Vectashield mounting media from Vector Laboratories Inc, Burlingame, CA, and imaged using a Leica SP8 upright confocal microscope (Zeiss) as described previously. All steps were performed at room temperature.

#### SVZ NSCs culture

The culture of SVZ NSCs was performed using SVZ wholemounts from postnatal (P12) C57BL/6J mice. The dissected tissues were placed in DMEM/F-12 (DF) medium containing 100 units/ml penicillin, 100 μg/ml streptomycin, and 250 ng/ml amphotericin B (abx). After pooling the tissues, they were incubated in 0.005% trypsin for 15 minutes at 37°C and pH 7.3. Subsequently, the tissues were transferred to uncoated T75 plastic tissue-culture dishes and kept overnight in N5 medium. The N5 medium consisted of DF supplemented with N2 supplements, 35 μg/ml bovine pituitary extract, abx, 5% FCS (HyClone), and 40 ng/ml EGF and basic FGF. Unattached cells were collected and plated again on uncoated plastic dishes. These cells were allowed to proliferate in N5 media until reaching 90-100% confluency, with fresh media added every other day for 3-4 days.

For experimentation, freshly cultured cells were always utilized. In **Fig. 2 A, C, E and F**, to study the activation of LV qNSCs, the cells were initially exposed to 10 ng/ml BMP4 for 24 hours to induce quiescence. Subsequently, carbachol treatment (15 μM) **(Fig. 2 A, C, E)** or a combination of carbachol (15 μM) and 4-DAMP (3 μM) **(Fig. 2 F)** was applied for 12 hours before the experiments were halted using 4% PFA, followed by staining as shown in the **Fig 2**.

In **Fig. 2 H**, after the freshly cultured cells were allowed to proliferate in N5 media until reaching 90-100% confluency, the growth factors and serum were removed from the culture media, and N6 media (containing N2 supplements, 35 μg/ml bovine pituitary extract, and antibiotics) were used to induce cell differentiation. As cell differentiation began, the cells in **Fig. 1 H** were treated daily with either carbachol alone (15 μM) or a combination of carbachol (15 μM) and 4-DAMP (3 μM). The experiments were halted after two days for immunostaining analysis, as described in further detail in **Fig. 2**.

#### SDS-PAGE and immunoblotting

Protein extracts were prepared from cultured SVZ NSCs, both carbachol-treated samples and control samples. The extracts were resolved by electrophoresis using SDS-PAGE and transferred onto nitrocellulose membranes. Antibodies were diluted in PBS containing 0.2% Triton X-100 (vol/vol) and 4% non-fat dry milk (wt/vol), followed by overnight incubation at 4 °C. Detection was achieved using secondary antibodies conjugated to horseradish peroxidase (#111-036-003, 106-006-003, 1:5 mL, Jackson ImmunoResearch) and treated with enhanced chemiluminescence (#1705061; Bio-Rad Laboratories). The intensity of DCX bands was quantified using ImageJ, and the values were then normalized against β-actin bands.

#### *In vitro* EdU staining

The SVZ NSCs cultures were plated on glass coverslips coated in DMEM/F-12. EdU staining was performed using Click-iT™ EdU Cell Proliferation Imaging Kit (ThermoFisher, USA) following the manufacturer’s protocol. EdU was added to the culture medium at concentration of 10 μM for 15 minutes. After labeling, the cells were washed three times with PBS. Next, the cells were permeabilized in 0.5% TBST for 30 seconds and fixed in 10% formaldehyde for 15 minutes. Subsequently, the cells were rinsed twice with PBS and incubated for 25 minutes in a Click-iT™ reaction cocktail contacting Click-iT™ reaction buffer, CuSO^4,^ Alexa fluor 647 Azide, and reaction buffer additive, while being protected from light. After staining, the cells on coverslips were washed three times with PBS containing 0.5% Triton X-100, 5 minutes each. Finally, the cells were counterstained with DAPI, mounted in vectashield mounting media (Vector Laboratories Inc, Burlingame, CA) and imaged using a Leica SP8 upright confocal microscope (Zeiss) as described above. All steps were performed at room temperature.

#### Total RNA extraction, cDNA synthesis, and Quantitative PCR

The SVZ NSCs cultures from control and carbachol-treated samples were extracted using the RNeasy plus Mini Kit (QIAGEN, USA), following the manufacturer’s protocol. Subsequently, first-strand cDNAs were synthesized using the SuperScript™ VILO™ cDNA Synthesis Kit (Invitrogen, USA) on a SimpliAmp™ thermocycler (ThermoFisher Scientific), as per the manufacturer’s instructions.

For the qPCR experiments, Power SYBR Green Master Mix (Applied Biosystems, USA) was used. The qPCR experiments were conducted on the StepOnePlus Real-Time PCR System running the StepOne Design and Analysis Software (Applied Biosystems). All reaction conditions and protocols were performed according to the manufacturer’s recommendations. The copies per reaction were determined by using absolute quantification.

The following primers were utilized for CHRM3, ITPR1, CAMK2D, MAPK10, and β-Actin genes:

ChRM3 F 5’ CCTCGCCTTTGTTTCCCAAC 3’

ChRM3 R 5’ TTGAGGAGAAATTCCCAGAGGT 3’

ITPR 1 F 5’ CGTTTTGAGTTTGAAGGCGTTT 3’

ITPR 1 R 5’ CATCTTGCGCCAATTCCCG 3’

CAMK2D F 5’ GATGGGGTAAAGGAGTCAACTG 3’

CAMK2D R 5’ CATTGTGGCATACAGCGACA 3’

MAPK10 F 5’ ACCACGAGCGGATGTCTTACT 3’

MAPK10 R 5’ GCACCCTACTGACCACAGG 3’

β-Actin F 5’ GGC TGT ATT CCC CTC CAT CG 3’

β-Actin R 5’ CCA GTT GGT AAC AAT GCC ATG 3’

#### Poly-A RNA Sequencing (RNAseq)

The total RNA extraction of the SVZ NSCs culture samples from total RNA extraction section were used to perform Poly-A-RNAseq experiments. In brief, SVZ NSCs cultures in proliferation media where the carbachol treatment was started on half of samples for 12 hours only as described in SVZ NSCs culture section. Then the total cellular RNA was extracted using RNeasy® Plus Mini Kit (Qiagen) and sent to LC Sciences company, USA. At LC Sciences, the RNA integrity of total RNA samples was checked using an Agilent Technologies 2100 Bioanalyzer. The poly-A containing mRNA molecules were purified using poly-T oligo attached magnetic beads and using two rounds of purification. After purification the poly-A RNA was fragmented using divalent cation containing buffer and elevated temperature. The first strand cDNA was obtained by reverse transcription using reverse transcriptase and a random primer. The second strand cDNA was synthesized by the process of removing the RNA template, synthesizing a replacement strand and dUTP method for strand-specificity. After the 3’-end adenylation and 5’-end phosphorylation, the blunt-ended cDNA adapters were ligated to the ends of the ds-cDNA. PCR amplification was performed to enrich the ligated cDNA and generate the library for sequencing. Quality control analysis and quantification of the DNA library were performed using agarose gel or Agilent Technologies 2100 Bioanalyzer High Sensitivity DNA Chip and KAPA Library Quantification Kits. Paired-end sequencing was performed on Illumina’s NovaSeq sequencing system.

#### Quantification and Statistical Analysis

All data are expressed as the difference between means ± standard error of the means (SEM). Statistical analyses were performed using GraphPad Prism (version 8) software. Paired t-tests were used for the analysis of *in vivo* optogenetics and chemogenetic stimulation and inhibition studies, as well as for the analysis of *in vitro* SVZ NSCs culture modulation studies and mRNA gene quantifications. Paired t-tests were also used for the analysis of subep-ChAT^+^ activity, as described in **SI Appendix, Fig. S1 and 3**. Furthermore, the activity of subep-ChAT^+^ neurons in **Fig. 3 and 4** was assessed in every batch of *in vivo* ACC-subep-ChAT^+^ circuit chemogenetic activation approach. Two-way ANOVA was used for the analysis of *in vitro* SVZ NSCs culture modulation and *in vivo* chemogenetic stimulation, as described in **Figs. 2 B, 3 G & P, and 4 E & M**.

A p-value of less than 0.05 was considered statistically significant. Detailed statistical results are described in figures and their respective legends. Significance levels were assigned as follows: * for p < 0.05, ** for p < 0.01, *** for p < 0.001 and **** for p < 0.0001.

To quantify the number of EdU^+^/GFAP^+^, CHRM3^+^/GFAP^+^,ITPR1^+^/GFAP^+^, CAMK2D^+^/GFAP^+^,MAPK10^+^/GFAP^+^, EdU^+^/Ki67^+^,CHRM3^+^/Ki67^+^, ITPR1^+^/Ki67^+^,CAMK2D^+^/Ki67^+^ and MAPK10^+^/Ki67^+^ cells in the SVZ wholemount, square images of 60μm × 60μm were captured in the V-SVZ and SVZ layers. Each image was centered on a subep-ChAT^+^ neuron, enabling the study of cell activation and proliferation adjacent to these neurons.

## Acknowledgments

We express our gratitude to Anne E West and Shawn Je for providing invaluable comments on the manuscript. We thank LC Sciences company for helping with Poly-A RNA sequencing experiment and analysis. This work was supported by NIH grant R01MH105416.

## Author Contributions

M.M.N. and H.H.Y. conceived the project and participated in research design. M.M.N. performed all experiments and analyzed data. M.M.N. and H.H.Y wrote the paper.

## Competing Interest Statement

The authors declare no competing financial interests.

## Notes

### Competing Interest Statement

The authors have declared no competing interest.

### Summary of Updates

The title of the manuscript.

## References

1. Lim DA & Alvarez-Buylla A (2016) The Adult Ventricular-Subventricular Zone (V-SVZ) and Olfactory Bulb (OB) Neurogenesis. Cold Spring Harb Perspect Biol 8(5).

2. Codega P, et al. (2014) Prospective identification and purification of quiescent adult neural stem cells from their in vivo niche. Neuron 82(3):545–559.

3. Mich JK, et al. (2014) Prospective identification of functionally distinct stem cells and neurosphere-initiating cells in adult mouse forebrain. Elife 3:e02669.

4. Doetsch F, Caille I, Lim DA, Garcia-Verdugo JM, & Alvarez-Buylla A (1999) Subventricular zone astrocytes are neural stem cells in the adult mammalian brain. Cell 97(6):703–716.

5. Obernier K, et al. (2018) Adult Neurogenesis Is Sustained by Symmetric Self-Renewal and Differentiation. Cell Stem Cell 22(2):221–234 e228.

6. Lois C, Garcia-Verdugo JM, & Alvarez-Buylla A (1996) Chain migration of neuronal precursors. Science 271(5251):978–981.

7. Ponti G, Obernier K, & Alvarez-Buylla A (2013) Lineage progression from stem cells to new neurons in the adult brain ventricular-subventricular zone. Cell Cycle 12(11):1649–1650.

8. Lois C & Alvarez-Buylla A (1994) Long-distance neuronal migration in the adult mammalian brain. Science 264(5162):1145–1148.

9. Luskin MB (1993) Restricted proliferation and migration of postnatally generated neurons derived from the forebrain subventricular zone. Neuron 11(1):173–189.

10. Petreanu L & Alvarez-Buylla A (2002) Maturation and death of adult-born olfactory bulb granule neurons: role of olfaction. J Neurosci 22(14):6106–6113.

11. Imayoshi I, et al. (2008) Roles of continuous neurogenesis in the structural and functional integrity of the adult forebrain. Nat Neurosci 11(10):1153–1161.

12. Livneh Y, Adam Y, & Mizrahi A (2014) Odor processing by adult-born neurons. Neuron 81(5):1097–1110.

13. Sakamoto M, et al. (2014) Continuous postnatal neurogenesis contributes to formation of the olfactory bulb neural circuits and flexible olfactory associative learning. J Neurosci 34(17):5788–5799.

14. Mak GK & Weiss S (2010) Paternal recognition of adult offspring mediated by newly generated CNS neurons. Nat Neurosci 13(6):753–758.

15. Sakamoto M, Kageyama R, & Imayoshi I (2014) The functional significance of newly born neurons integrated into olfactory bulb circuits. Front Neurosci 8:121.

16. Soudry Y, Lemogne C, Malinvaud D, Consoli SM, & Bonfils P (2011) Olfactory system and emotion: common substrates. Eur Ann Otorhinolaryngol Head Neck Dis 128(1):18–23.

17. Ihrie RA & Alvarez-Buylla A (2011) Lake-front property: a unique germinal niche by the lateral ventricles of the adult brain. Neuron 70(4):674–686.

18. Lazarini F & Lledo PM (2011) Is adult neurogenesis essential for olfaction? Trends Neurosci 34(1):20–30.

19. Robel S, Berninger B, & Gotz M (2011) The stem cell potential of glia: lessons from reactive gliosis. Nat Rev Neurosci 12(2):88–104.

20. Benner EJ, et al. (2013) Protective astrogenesis from the SVZ niche after injury is controlled by Notch modulator Thbs4. Nature 497(7449):369–373.

21. Jones KS & Connor B (2012) Intrinsic regulation of adult subventricular zone neural progenitor cells and the effect of brain injury. Am J Stem Cells 1(1):48–58.

22. Obernier K & Alvarez-Buylla A (2019) Neural stem cells: origin, heterogeneity and regulation in the adult mammalian brain. Development 146(4).

23. Berg DA, Belnoue L, Song H, & Simon A (2013) Neurotransmitter-mediated control of neurogenesis in the adult vertebrate brain. Development 140(12):2548–2561.

24. Bovetti S, Gribaudo S, Puche AC, De Marchis S, & Fasolo A (2011) From progenitors to integrated neurons: role of neurotransmitters in adult olfactory neurogenesis. J Chem Neuroanat 42(4):304–316.

25. Young SZ, Taylor MM, & Bordey A (2011) Neurotransmitters couple brain activity to subventricular zone neurogenesis. Eur J Neurosci 33(6):1123–1132.

26. Paez-Gonzalez P, Asrican B, Rodriguez E, & Kuo CT (2014) Identification of distinct ChAT(+) neurons and activity-dependent control of postnatal SVZ neurogenesis. Nat Neurosci 17(7):934–942.

27. Naffaa MM, Khan RR, Kuo CT, & Yin HH (2023) Cortical regulation of neurogenesis and cell proliferation in the ventral subventricular zone. Cell Rep 42(7):112783.

28. Luo J, Busillo JM, & Benovic JL (2008) M3 muscarinic acetylcholine receptor-mediated signaling is regulated by distinct mechanisms. Mol Pharmacol 74(2):338–347.

29. Ambudkar IS (2016) Calcium signalling in salivary gland physiology and dysfunction. J Physiol 594(11):2813–2824.

30. Arige V, et al. (2022) Functional determination of calcium-binding sites required for the activation of inositol 1,4,5-trisphosphate receptors. Proc Natl Acad Sci U S A 119(39):e2209267119.

31. Wayman GA, Lee YS, Tokumitsu H, Silva AJ, & Soderling TR (2008) Calmodulin-kinases: modulators of neuronal development and plasticity. Neuron 59(6):914–931.

32. Najar MA, et al. (2021) A complete map of the Calcium/calmodulin-dependent protein kinase kinase 2 (CAMKK2) signaling pathway. J Cell Commun Signal 15(2):283–290.

33. Hampel H, et al. (2018) The cholinergic system in the pathophysiology and treatment of Alzheimer’s disease. Brain 141(7):1917–1933.

34. Poulin B, et al. (2010) The M3-muscarinic receptor regulates learning and memory in a receptor phosphorylation/arrestin-dependent manner. Proc Natl Acad Sci U S A 107(20):9440–9445.

35. Felder CC (1995) Muscarinic acetylcholine receptors: signal transduction through multiple effectors. FASEB J 9(8):619–625.

36. Caulfield MP (1993) Muscarinic receptors--characterization, coupling and function. Pharmacol Ther 58(3):319–379.

37. Liang J, et al. (2010) Type 3 inositol 1,4,5-trisphosphate receptor negatively regulates apoptosis during mouse embryonic stem cell differentiation. Cell Death Differ 17(7):1141–1154.

38. Patergnani S, et al. (2020) Various Aspects of Calcium Signaling in the Regulation of Apoptosis, Autophagy, Cell Proliferation, and Cancer. Int J Mol Sci 21(21).

39. Hook SS & Means AR (2001) Ca(2+)/CaM-dependent kinases: from activation to function. Annu Rev Pharmacol Toxicol 41:471–505.

40. Zalcman G, Federman N, & Romano A (2018) CaMKII Isoforms in Learning and Memory: Localization and Function. Front Mol Neurosci 11:445.

41. Lu DZ, et al. (2020) CaMKII(delta) regulates osteoclastogenesis through ERK, JNK, and p38 MAPKs and CREB signalling pathway. Mol Cell Endocrinol 508:110791.

42. Si J & Collins SJ (2008) Activated Ca2+/calmodulin-dependent protein kinase IIgamma is a critical regulator of myeloid leukemia cell proliferation. Cancer Res 68(10):3733–3742.

43. Giovannini MG, et al. (2001) Mitogen-activated protein kinase regulates early phosphorylation and delayed expression of Ca2+/calmodulin-dependent protein kinase II in long-term potentiation. J Neurosci 21(18):7053–7062.

44. Semba T, et al. (2020) JNK Signaling in Stem Cell Self-Renewal and Differentiation. Int J Mol Sci 21(7).

45. Bengoa-Vergniory N, Gorrono-Etxebarria I, Gonzalez-Salazar I, & Kypta RM (2014) A switch from canonical to noncanonical Wnt signaling mediates early differentiation of human neural stem cells. Stem Cells 32(12):3196–3208.

46. Cargnello M & Roux PP (2011) Activation and function of the MAPKs and their substrates, the MAPK-activated protein kinases. Microbiol Mol Biol Rev 75(1):50–83.

47. Sanai N, et al. (2011) Corridors of migrating neurons in the human brain and their decline during infancy. Nature 478(7369):382–386.

