## Supplemental figures for "A Cholinergic Signaling Pathway underlying Cortical Circuit Regulation of Lateral Ventricle Quiescent Neural Stem Cells"

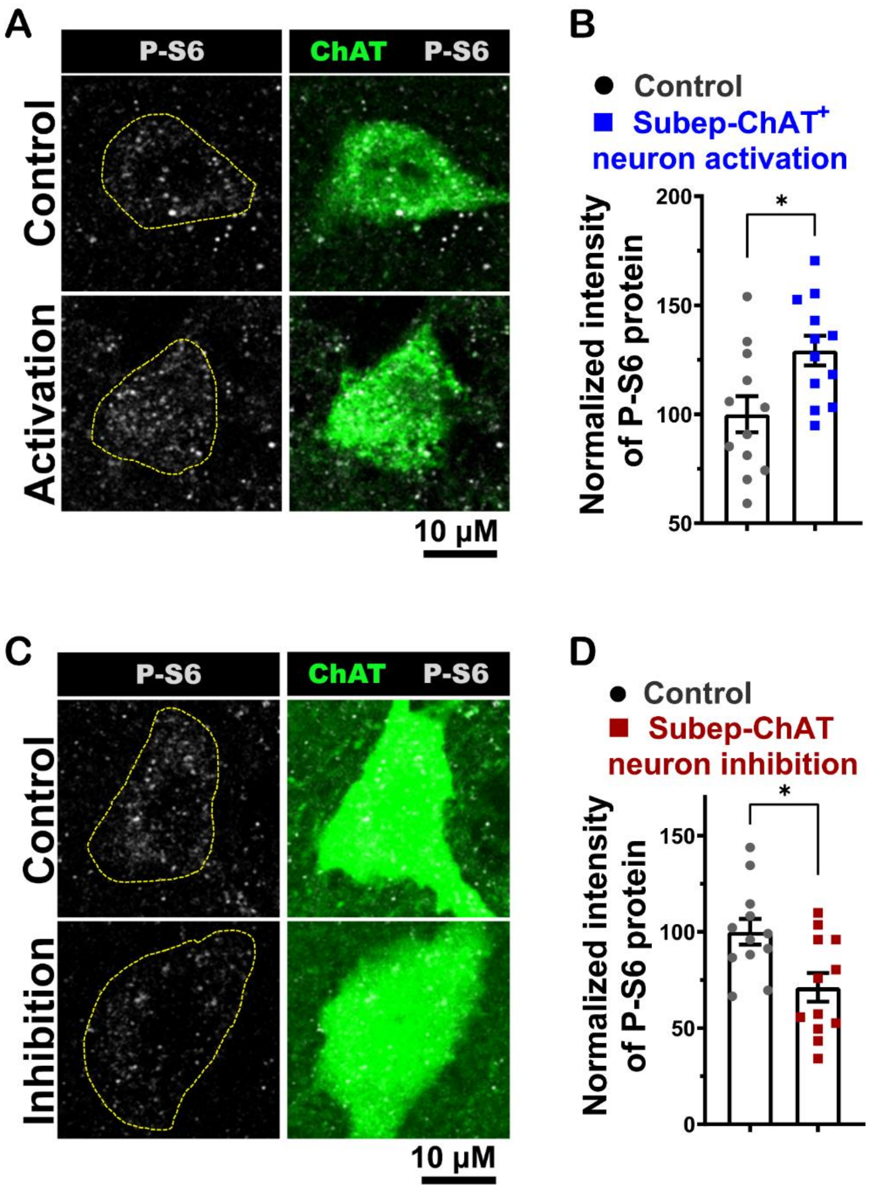


**SI Appendix, Fig. S1. Assessment of subep-ChAT⁺ neuron activity post *in vivo* optogenetic activation and inhibition. (A)** ChAT (green) and P-S6 (gray) immunofluorescence staining of subep-ChAT⁺ neuron in control SVZ wholemount (upper images) vs. activation SVZ wholemount (lower images) from stimulated mice in **Fig. 1 A. (B)** P-S6 intensity analysis of subep-ChAT⁺ neurons in activation vs. control SVZ wholemounts. $P < 0.031$, $t_{11} = 2.5$, n=12, Paired t-test. Data collected from four stimulated *Cr-Cre* mice. Each dot represents a subep-ChAT⁺ neuron. **(C)** ChAT (green) and P-S6 (gray) immunofluorescence staining of subep-ChAT⁺ neuron in control SVZ wholemount (upper images) vs. inhibition SVZ wholemount (lower images) from stimulated mice in **Fig. 1 E. (D)** P-S6 intensity analysis of subep-ChAT⁺ neurons in control vs. inhibition SVZ wholemounts. $P < 0.0332$, $t_{11} = 2.4$, n=12, Paired t-test. Data collected from four stimulated *Cr-Cre* mice. Each dot represents a subep-ChAT⁺ neuron. All error bars indicate SEM.

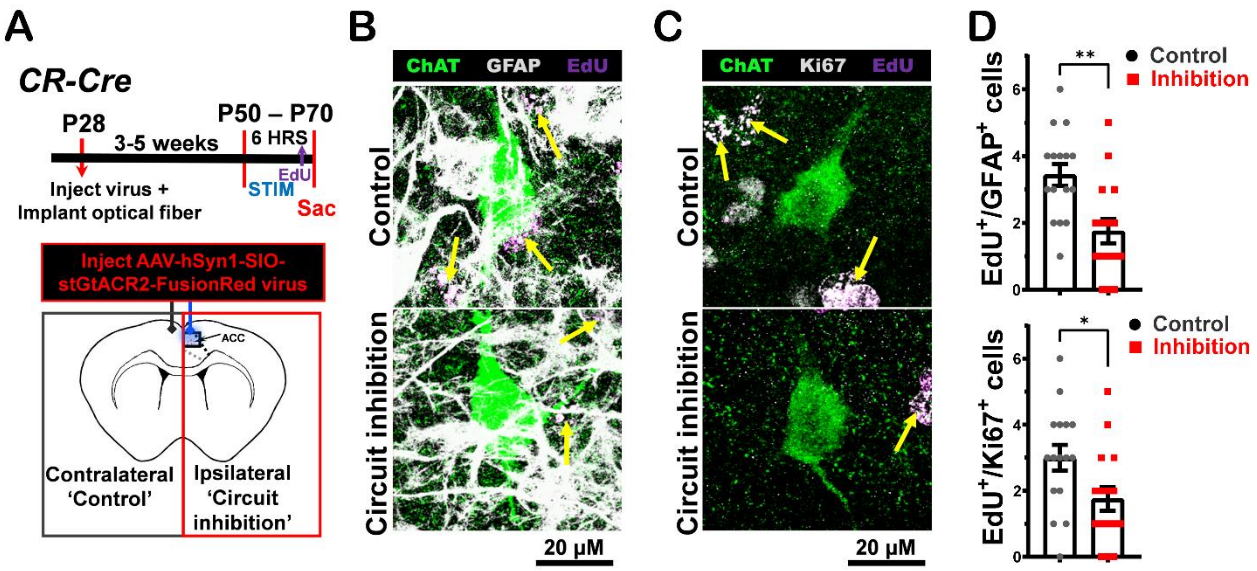


**SI Appendix, Fig. S2. *In vivo* regulation of LV NSCs via optogenetic inhibition of ACC-subep-ChAT⁺ circuit. (A)** Experimental design (upper) and schematic representation (lower) of *in vivo* optogenetic inhibition for 6hrs post AAV-hSyn1-SIO-stGtACR2-FusionRed virus injection and optical fibers implantation into ACC regions of (P28) *CR-Cre* mice. **(B)** Immunofluorescence staining for ChAT (green), GFAP (gray) and EdU (purple) of ipsilateral SVZ wholemount (lower images; ACC-subep-$ChAT^+$ circuit inhibition) vs. contralateral SVZ wholemount (upper images; (control)) from mice in panel A. Yellow arrows show $EdU^+$/$GFAP^+$ cells surrounding subep-$ChAT^+$ neurons. **(C)** Immunofluorescence staining for ChAT (green), Ki67 (gray) and EdU (purple) of ipsilateral SVZ wholemount (lower images; ACC-subep-$ChAT^+$ circuit inhibition) vs. contralateral SVZ wholemount (upper images; (control)) from mice in panel A. Yellow arrows show $EdU^+$/$Ki67^+$ cells surrounding subep-$ChAT^+$ neurons. **(D)** Upper; analysis of $EdU^+$/$GFAP^+$ cells per a subep-$ChAT^+$ neurons in SVZ wholemounts of ipsilateral vs. contralateral in panel F. $P < 0.0018$, $t_{15} = 3.8$, n=16, Paired t-test. Data collected from four stimulated *CR-Cre* mice. Each dot represents total $EdU^+$/$GFAP^+$ cells surrounding a subep-$ChAT^+$ neuron. Lower; analysis of $EdU^+$/$Ki67^+$ cells per a subep-$ChAT^+$ neurons in SVZ wholemounts of ipsilateral vs. contralateral in panel G. $P < 0.041$, $t_{15} = 2.24$, n=16, Paired t-test. Data collected from four stimulated *CR-Cre* mice. Each dot represents total $EdU^+$/$Ki67^+$ cells surrounding a subep-$ChAT^+$ neuron. All error bars indicate SEM.

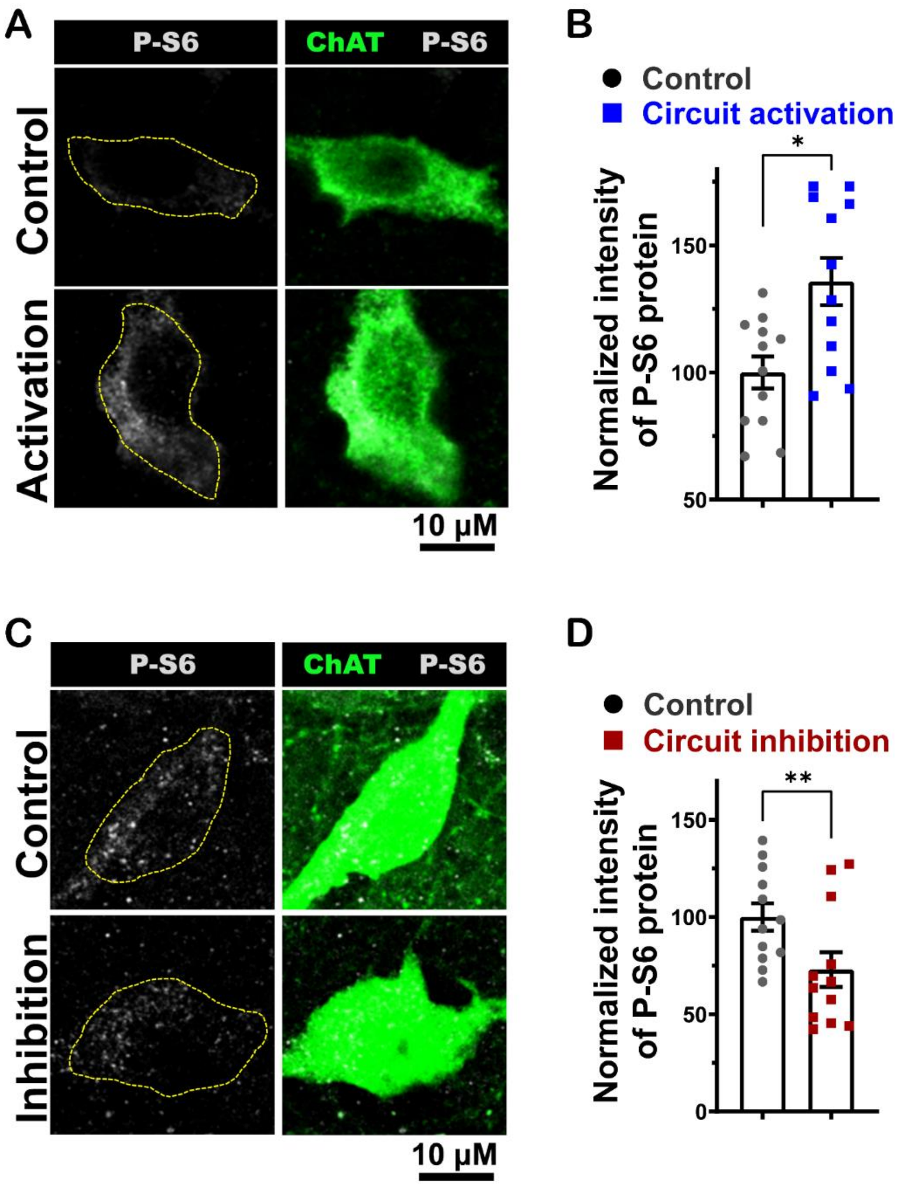


**SI Appendix, Fig. S3. Assessment of subep-ChAT$^+$ neuron activity post *in vivo* chemogenetic and optogenetic ACC-subep-ChAT$^+$ circuit activation and inhibition, respectively. (A)** ChAT (green) and P-S6 (gray) immunofluorescence staining of subep-ChAT$^+$ neuron in control SVZ wholemount (upper images) vs. ACC-subep-ChAT$^+$ circuit activation SVZ wholemount (lower images) from stimulated mice in **Fig. 1 I. (B)** P-S6 intensity analysis of subep-ChAT$^+$ neurons in activation vs. control SVZ wholemounts. $P < 0.0162$, $t_{11} = 2.84$, n=12, Paired t-test. Data collected from four stimulated *Cr-Cre* mice. Each dot represents a subep-ChAT$^+$ neuron. **(C)** ChAT (green) and P-S6 (gray) immunofluorescence staining of subep-ChAT$^+$ neuron in control SVZ wholemount (upper images) vs. ACC-subep-ChAT$^+$ circuit inhibition SVZ wholemount (lower images) from stimulated mice in **SI Appendix, Fig. 3A. (D)** P-S6 intensity analysis of subep-ChAT$^+$ neurons in control vs. inhibition SVZ wholemounts. $P < 0.0070$, $t_{11} = 3.3$, n=12, Paired t-test. Data collected from four stimulated *Cr-Cre* mice. Each dot represents a subep-ChAT$^+$ neuron. All error bars indicate SEM.

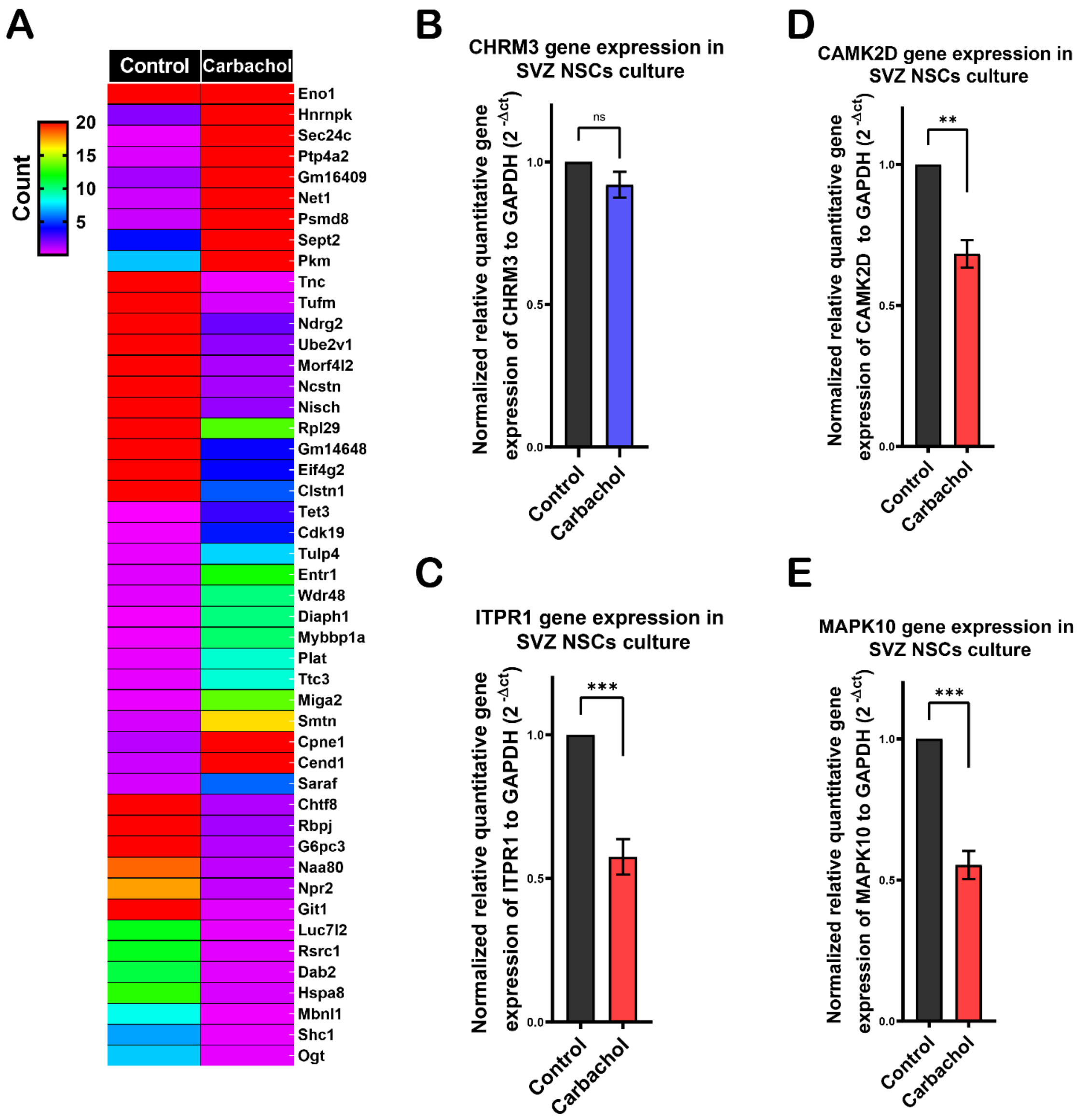


**Appendix, Fig. S4. Bulk RNA-Seq analysis and RT-qPCR results for CHRM3, ITPR1, CAMK2D and MAPK10. (A)** Heat map illustrating RNA-Seq of representative differential expression transcripts induced by treating SVZ NSCs culture with carbachol (15μM) versus control in proliferation media collected after 12 hrs. **(B)** Normalized relative quantitative gene expression of ChRM3/β-actin from RT-qPCR experiment in control versus treatment samples with carbachol (15μM) of SVZ NSCs cultures in proliferation media collected after 12 hrs. **(C)** Normalized relative quantitative gene expression of ITPR1/β-actin from RT-qPCR experiment in control versus treatment samples with carbachol (15μM) of SVZ NSCs cultures in proliferation media collected after 12 hrs. **(D)** Normalized relative quantitative gene expression of CAMK2D/β-actin from RT-qPCR experiment in control versus treatment samples with carbachol (15μM) of SVZ NSCs cultures in proliferation media collected after 12 hrs. **(E)** Normalized relative quantitative gene expression of MAPK10/β-actin from RT-qPCR experiment in control versus treatment samples with carbachol (15μM) of SVZ NSCs cultures in proliferation media collected after 12 hrs.

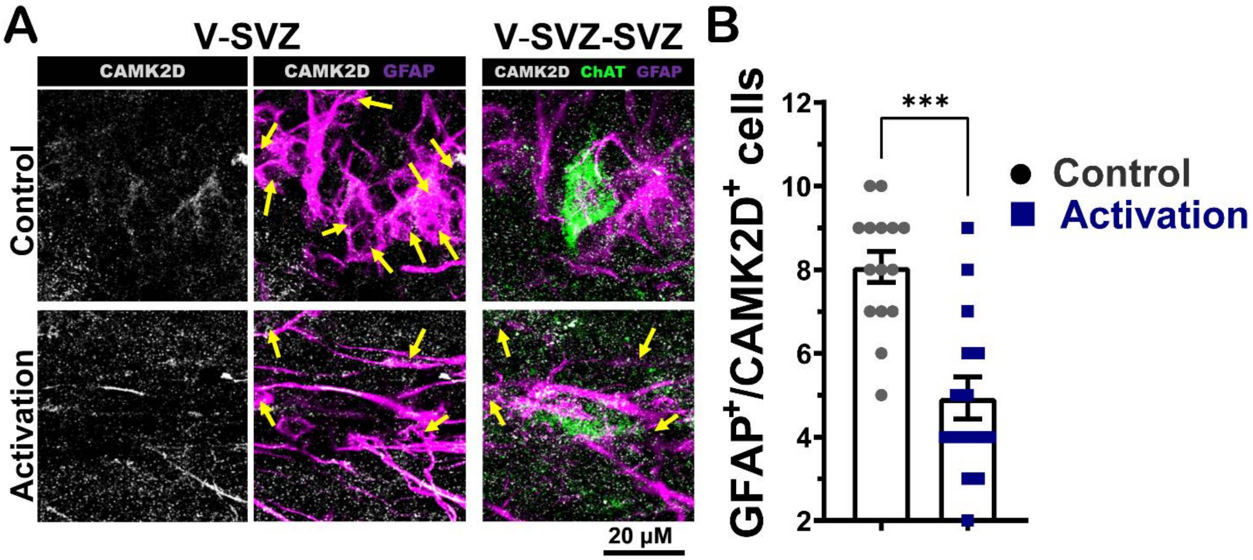


**SI Appendix, Fig. S5. *In vivo* CAMK2D activation control LV NSCs proliferative activity.** **(A)** Immunofluorescence staining for CAMK2D (gray), ChAT (green), and GFAP (purple) of ipsilateral SVZ wholemount (lower images; ACC-subep-ChAT$^+$ circuit activation) vs. contralateral SVZ wholemount (upper images; (control)) from mice in **Fig. 3C**. Yellow arrows show GFAP$^+$/ CAMK2D$^+$ cells surrounding a subep-ChAT$^+$ neuron. **(B)** Analysis of GFAP$^+$/CAMK2D$^+$ cells per a subep-ChAT$^+$ neuron in SVZ wholemounts of ipsilateral vs. contralateral in panel C. $P < 0.0002$, $t_{14} = 4.9$, n=15, Paired t-test. Data collected from three stimulated *CR-Cre* mice. Each dot represents total GFAP$^+$/ CAMK2D$^+$ cells surrounding a subep-ChAT$^+$ neuron. All error bars indicate SEM.

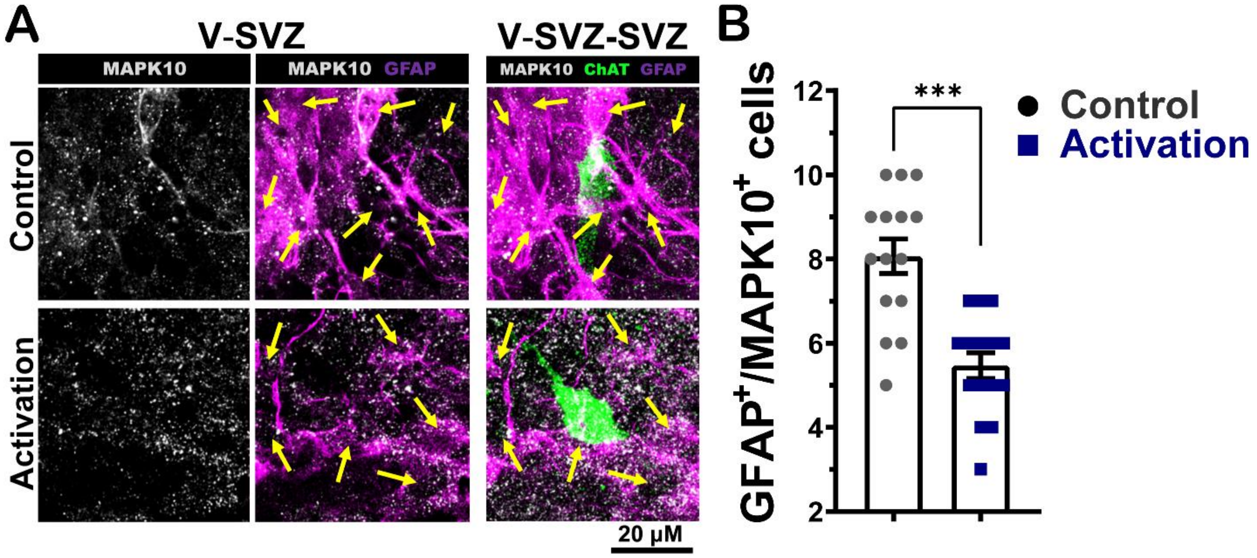


**SI Appendix, Fig. S6.** ***In vivo*** **MAPK10 activation control LV NSCs proliferative activity.** **(A)** Immunofluorescence staining for MAPK10 (gray), ChAT (green), and GFAP (purple) of ipsilateral SVZ wholemount (lower images; ACC-subep-ChAT$^+$ circuit activation) vs. contralateral SVZ wholemount (upper images; (control)) from mice in **Fig. 3C**. Yellow arrows show GFAP$^+$/ MAPK10$^+$ cells surrounding a subep-ChAT$^+$ neuron. **(B)** Analysis of GFAP$^+$/ MAPK10$^+$ cells per a subep-ChAT$^+$ neuron in SVZ wholemounts of ipsilateral vs. contralateral in panel C. $P < 0.0002$, $t_{14} = 4.9$, n=15, Paired t-test. Data collected from three stimulated *CR-Cre* mice. Each dot represents total GFAP$^+$/ MAPK10$^+$ cells surrounding a subep-ChAT$^+$ neuron. All error bars indicate SEM.
